# Membrane Hydrophobicity Determines the Activation Free Energy of Passive Lipid Transport

**DOI:** 10.1101/2021.03.17.435885

**Authors:** Julia R. Rogers, Gustavo Espinoza Garcia, Phillip L. Geissler

## Abstract

The collective behavior of lipids with diverse chemical and physical features determines a membrane’s thermodynamic properties. Yet, the influence of lipid physicochemical properties on lipid dynamics, in particular interbilayer transport, remains underexplored. Here, we systematically investigate how the activation free energy of passive lipid transport depends on lipid chemistry and membrane phase. Through all-atom molecular dynamics simulations of 11 chemically distinct glycerophospholipids, we determine how lipid acyl chain length, unsaturation, and headgroup influence the free energy barriers for two elementary steps of lipid transport, lipid desorption, which is rate-limiting, and lipid insertion into a membrane. Consistent with previous experimental measurements, we find that lipids with longer, saturated acyl chains have increased activation free energies compared to lipids with shorter, unsaturated chains. Lipids with different headgroups exhibit a range of activation free energies; however, no clear trend based solely on chemical structure can be identified, mirroring difficulties in the interpretation of previous experimental results. Compared to liquid-crystalline phase membranes, gel phase membranes exhibit substantially increased free energy barriers. Overall, we find that the activation free energy depends on a lipid’s local hydrophobic environment in a membrane and that the free energy barrier for lipid insertion depends on a membrane’s interfacial hydrophobicity. Both of these properties can be altered through changes in lipid acyl chain length, lipid headgroup, and membrane phase. Thus, the rate of lipid transport can be tuned through subtle changes in local membrane composition and order, suggesting an unappreciated role for nanoscale membrane domains in regulating cellular lipid dynamics.

**SIGNIFICANCE:** Cell homeostasis requires spatiotemporal regulation of heterogeneous membrane compositions, in part, through non-vesicular transport of individual lipids between membranes. By systematically investigating how the chemical diversity present in glycerophospholipidomes and variations in membrane order influence the free energy barriers for passive lipid transport, we discover a correlation between the activation free energy and membrane hydrophobicity. By demonstrating how membrane hydrophobicity is modulated by local changes in membrane composition and order, we solidify the link between membrane physicochemical properties and lipid transport rates. Our results suggest that variations in cell membrane hydrophobicity may be exploited to direct non-vesicular lipid traffic.

## INTRODUCTION

Over a thousand chemically diverse lipid species are heterogeneously distributed among eukaryotic cell membranes (1, 2). Even within a single membrane, compositional differences exist between leaflets (3) and laterally between nanodomains (4). Because the collective behavior of a membrane’s constituent lipids determines its physical properties, such as fluidity, thickness, and curvature, membrane compositions are under homeostatic control (5–7). One way that proper lipid distributions are maintained is through non-vesicular transport of individual lipids between membranes. Non-vesicular transport enables rapid and specific alteration of membrane compositions, such as required to withstand cellular stress (8, 9).

Despite the recognized importance of lipid chemistry in determining membrane physical properties, how lipid physicochemical properties influence the dynamic processes that maintain precise membrane compositions is poorly understood. *In vivo*, lipid transfer proteins may be largely responsible for selectively transferring lipids recognized through specific protein–lipid interactions. Lipid transfer proteins are also equipped with membrane binding domains or motifs that may enable them to target donor and acceptor membranes with particular compositions and biophysical properties (10, 11). As demonstrated by *in vitro* experiments, lipids with subtle chemical differences are also passively exchanged between membranes at different rates (12–25). For example, diacyl phosphatidylcholine (PC) exchanges much more slowly than lysoPC, its single-tailed counterpart (14, 20). Even the addition of just two carbons to an acyl chain of a phospholipid reduces its exchange rate by roughly 10-fold (14–23). The physical properties and chemical composition of the donor and acceptor membranes additionally influence the rate at which a lipid is passively transported (19–26). For example, dimristoylphosphatidylcholine (DMPC) exchanges more rapidly between liquid-crystalline (L_*α*_) phase vesicles than more ordered gel (L_*β*_) phase ones (20, 26). Therefore, the underlying free energy barriers for transport also depend on a lipid’s chemical structure and properties of the donor and acceptor membranes.

During passive lipid transport, a lipid first desorbs from a donor membrane, diffuses through solvent, and then inserts into an acceptor membrane. Due to the large free energetic cost of disrupting a lipid’s local hydrophobic environment in a membrane, lipid desorption is the rate-limiting step and, thus, determines the activation free energy of lipid transport. A smaller free energy barrier exists for lipid insertion (13, 18, 20, 23, 27). We recently discovered the molecular origins of both free energy barriers by identifying the reaction coordinate for passive DMPC transport in unbiased molecular dynamics (MD) simulations. The reaction coordinate measures the extent of hydrophobic contact between the transferring DMPC and membrane. Importantly, we demonstrated that the reaction coordinate accurately captures the dynamics of DMPC transport observed in unbiased all-atom MD simulations, locates transition states with high fidelity, and, thus, resolves the rate-limiting free energy barriers for DMPC desorption and insertion. The free energy barrier for DMPC desorption reflects the thermodynamic cost of breaking hydrophobic lipid–membrane contacts, and the free energy barrier for DMPC insertion reflects the cost of disrupting the membrane–solvent interface during the formation of initial hydrophobic lipid–membrane contacts (27).

Here, we systematically investigate how the free energy barriers for both lipid desorption and insertion vary with lipid chemistry and membrane phase. By focusing our analysis on glycerophospholipids, we determine how the chemical diversity found among structural lipids in eukaryotic lipidomes (1, 2) influences the rate of passive lipid transport. Since all glycerophospholipids possess the same general physical properties, we assume at the outset that the reaction coordinate for DMPC transport discovered in our previous work (27) also captures the salient features of transporting other glycerophospholipids. Working under this assumption allows us to efficiently quantify the free energy barriers for transporting lipids between 14 different membranes using all-atom MD simulations. The results reported herein indicate that the biophysical mechanism of lipid transport is indeed invariant to glycerophospholipid chemistry, supporting our initial assumption. We find that the activation free energy for lipid transport increases as lipid acyl chain length increases and as membrane order increases such that L_*β*_ phase membranes pose the largest barriers for lipid transport. Additionally, we find the activation free energy to be strongly dependent on the identity of the lipid headgroups. Our results are consistent with *in vitro* measurements and provide a biophysical rationale for previously unexplained experimental observations. Furthermore, the atomistic detail provided by molecular simulations allows us to correlate the free energy barriers for lipid desorption and insertion with the molecular properties of a membrane and, thus, to identify the key physicochemical properties that control transport rates: Membrane hydrophobicity, specifically the number of hydrophobic contacts a lipid makes with surrounding membrane lipids, determines the rate of passive lipid transport.

## METHODS

The rate-limiting step of passive lipid transport – spontaneous desorption of a lipid from a membrane – occurs over minutes to hours. As a result, calculating the transport rate directly from unbiased MD simulations is computationally intractable for even one membrane system. Instead, our approach to quantitatively assess how the rate of passive lipid transport varies with lipid chemistry and membrane phase capitalizes on knowledge of the reaction coordinate (27), which we assume to be the same for all lipids investigated.

Free energy profiles along order parameters other than the reaction coordinate generally yield barriers that underestimate the rate-determining activation free energy, and therefore do not provide sufficient thermodynamic information to accurately determine barrier-crossing rates (28). A barrier can even be entirely absent from such a free energy profile, in which case even a qualitative inference of the transition dynamics is erroneous. As we have shown previously, lipid insertion is a prime example of this pathology: Free energy profiles calculated as a function of the center of mass displacement of the lipid from the membrane incorrectly suggest that lipid insertion is a barrier-less process (27).

By calculating free energy profiles along the reaction coordinate, we efficiently determined the rate-limiting free energy barrier for transporting 11 different lipids between L_*α*_ phase membranes composed of the same lipid species as the one being transported. For high melting temperature lipids, free energy barriers for lipid desorption from and insertion into both L_*α*_ and L_*β*_ phase membranes were determined (Fig. 1).

**Figure 1:**
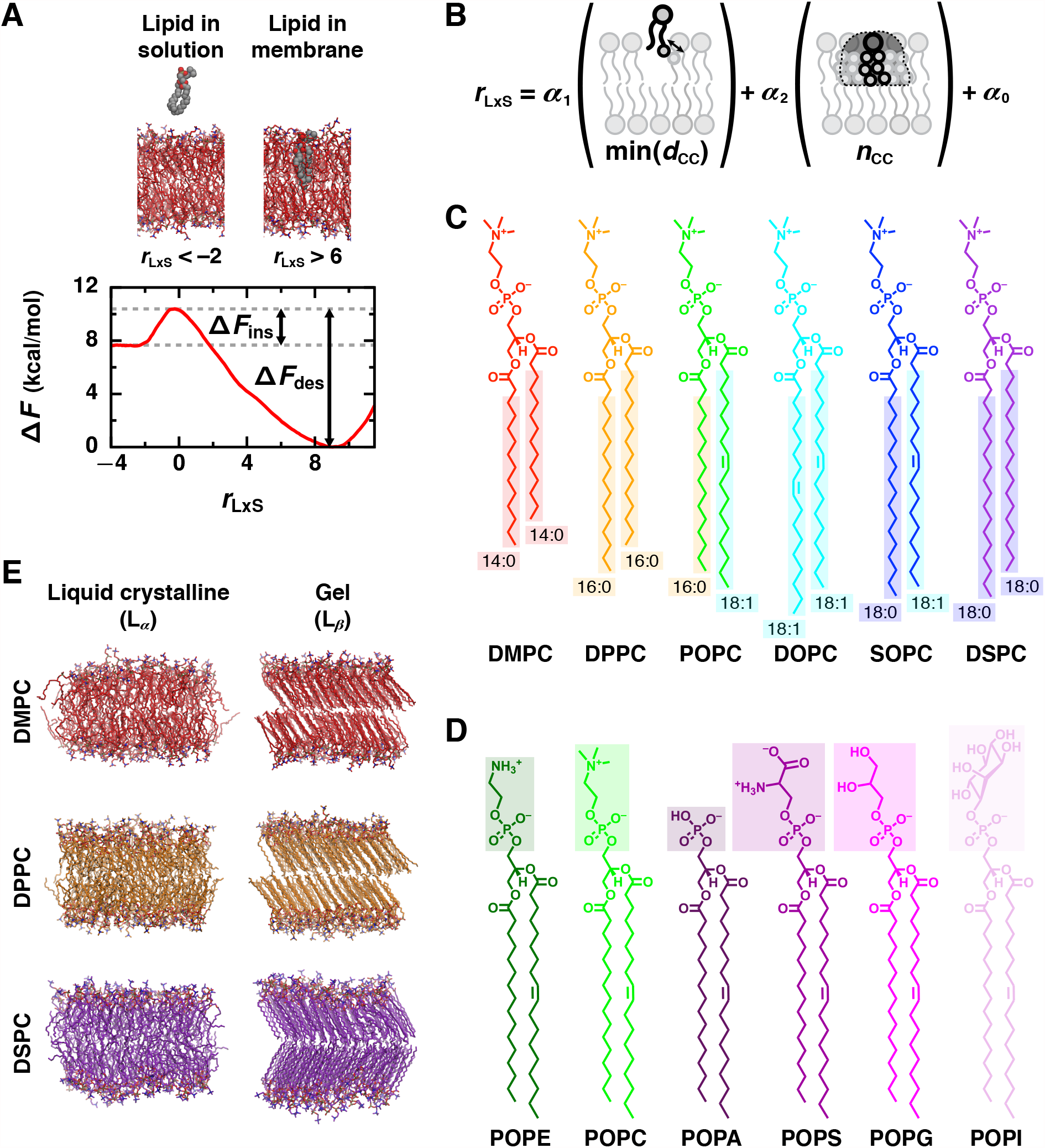
Physicochemical properties that determine the rate of lipid transport were assessed for a variety of lipid chemistries and phases by calculating the free energy barriers for lipid desorption and insertion. (A) An illustrative free energy profile, calculated as a function of the reaction coordinate, *r*_LxS_, for a L_*α*_ phase DMPC membrane, reveals barriers for lipid desorption (Δ *F*_des_) and lipid insertion (Δ *F*_ins_). Representative configurations are shown for negative values of *r*_LxS_ (where the lipid is in solution) and for positive values of *r*_LxS_ (where the lipid is in the membrane). Solvent is not rendered. (B) *r*_LxS_ is a linear combination of min *d*_CC_ (the minimum distance between any hydrophobic carbon of the lipid and of the closest membrane leaflet) and *n*_CC_ (the number of close hydrophobic carbon contacts between the lipid and closest leaflet). (C and D) Chemical structures of the lipids investigated with varied (C) acyl chain lengths and degrees of saturation, and (D) headgroups. POPC’s structure is shown in both (C) and (D). In (C), hydrophobic carbons are boxed. (E) Representative configurations of membranes composed of DMPC, DPPC, or DSPC in both L_*α*_ and L_*β*_ phases.

### Simulated systems

To systematically assess how the activation free energy of lipid transport depends on lipid physicochemical properties, we simulated membrane systems composed of one of 11 different glycerophospholipid species. This included a series of PC lipids with increasing acyl chain lengths and various degrees of saturation (Fig. 1C): 1,2-dimyristoyl-*sn*-glycero-3-PC (DMPC), 1,2-dipalmitoyl-*sn*-glycero-3-PC (DPPC), 1-palmitoyl-2-oleoyl-*sn*-glycero-3-PC (POPC), 1,2-dioleoyl-*sn*-glycero-3-PC (DOPC), 1-stearoyl-2-oleoyl-*sn*-glycero-3-PC (SOPC), and 1,2-distearoyl-*sn*-glycero-3-PC (DSPC). This also included a series of 1-palmitoyl-2-oleoyl (PO) lipids with both zwitterionic and anionic headgroups (Fig. 1D): PO-*sn*-glycero-3-phosphoethanolamine (POPE), POPC, PO-*sn*-glycero-3-phosphate (POPA), PO-*sn*-glycero-3-phospho-L-serine (POPS), PO-*sn*-glycero-3-phospho-(1’-*rac*-glycerol) (POPG), and PO-*sn*-glycero-3-phosphoinositol (POPI). Membranes composed of each lipid were simulated in the L_*α*_ phase at 320K with the exception of DSPC, which was simulated at 350K due to its higher melting temperature. Membranes composed of the saturated lipids, DMPC, DPPC, and DSPC, were additionally simulated in the L_*β*_ phase (Fig. 1E) at 275K, 295K, and 320K, respectively.

For all lipid species, initial L_*α*_ phase bilayers of 128 lipids (64 per leaflet) surrounded by 3.2 nm thick slabs of solvent were built using the CHARMM-GUI Membrane Builder (29, 30). We expect results reported herein to be invariant to our choice of membrane size since similar free energy calculations have been shown to be membrane size-independent (31). Neutralizing sodium ions were added to each anionic membrane. The L_*α*_ phase DSPC bilayer was also used to initialize a simulation at 320K that yielded a L_*β*_ phase bilayer within 100 ns. Configurations of L_*β*_ phase DPPC and DMPC bilayers were instead used to initialize simulations at temperatures consistent with a L_*β*_ phase to avoid these systems getting trapped in the ripple phase upon cooling, which can occur in simulations of shorter chain lipids (32). The initial L_*β*_ phase DPPC bilayer was obtained from the LipidBook repository (33) and also used to construct an initial L_*β*_ phase DMPC bilayer. The CHARMM36 force field (34) was used to model all lipids in combination with the CHARMM TIP3P water model (35).

### Molecular dynamics simulations

For each different lipid species and membrane phase, solvated bilayers were simulated to characterize the physical properties of each membrane at equilibrium. All simulations were performed in an isothermal–isobaric (NPT) ensemble using GROMACS 2019 (36). The pressure was maintained at 1 bar using semi-isotropic pressure coupling with an isothermal compressibility of 4.5 ×10 −5 bar^−1^, and the temperature was maintained using the Nosé-Hoover thermostat (37, 38) with a coupling time constant of 1 ps. The lipids and solvent were coupled to separate thermostats. Dynamics were evolved using the leapfrog algorithm (39) and a 2 fs time step. All bonds to hydrogen were constrained using the LINCS algorithm (40). Lennard-Jones forces were smoothly switched off between 0.8 and 1.2 nm. Coulomb interactions were truncated at 1.2 nm, and long-ranged Coulomb interactions were calculated using Particle Mesh Ewald (PME) summation (41). Neighbor lists were constructed with the Verlet list cut-off scheme (42).

Each initial bilayer configuration was first energy minimized using the steepest descent algorithm and then equilibrated in two steps: The first 250 ps equilibration used the Berendsen barostat (43) to maintain the pressure with a coupling time constant of 2 ps, and the second 250 ps equilibration used the Parinello-Rahman barostat (44) with a coupling time constant of 5 ps. To allow the bilayers’ structures to fully equilibrate, runs of 50 ns for the L_*α*_ and of 100 ns for the L_*β*_ phase membranes were performed using the same parameters as the second equilibration step. The final configurations from these runs were used to construct initial configurations for enhanced sampling simulations that had an additional lipid free in solution. Unbiased simulations of each bilayer were run for an additional 300 ns, which was analyzed to calculate average properties of each membrane.

### Free energy calculations

To determine the activation free energy for passively transporting a lipid between membranes, we calculated free energy profiles as a function of the reaction coordinate (Fig. 1A). As determined in our previous study (27), the reaction coordinate, *r*_LxS_ (which was labeled *r*_*c*_ in (27)), is a linear combination of two order parameters that measure hydrophobic lipid–membrane contacts between the transferring lipid and closest membrane leaflet (Fig. 1B). Both parameters are based on the collection of distances 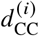 between a hydrophobic carbon of the lipid and a hydrophobic carbon of the membrane, where *i* labels a particular carbon-carbon (CC) pair: (1) 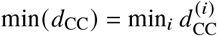 is the minimum of these CC distances; and (2) *n*_CC_ is the number of CC pairs that satisfy 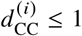 nm. This cutoff value includes roughly two carbon solvation shells around a hydrophobic carbon of a lipid in a bilayer. The hydrophobic carbons that are considered to calculate *n*_CC_ for each lipid are boxed in Fig. 1C. In detail,

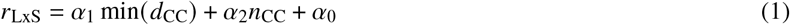

with coefficients *α*_1_ = − 2.247 nm^−1^, *α*_2_ = 0.004828, and *α*_0_ = 0.6014 such that *r*_LxS_ is a unitless quantity. By construction, *r*_LxS_ = 0 at the free energy barrier that separates configurations with the transferring lipid fully in solution, which have negative values of *r*_LxS_, and configurations with the transferring lipid fully in the membrane, which have positive values of *r*_LxS_ (Fig. 1A). Regardless of its total number of hydrophobic carbons, a lipid forms only a few hydrophobic contacts with the membrane at the free energy barrier (27). The largest free energy barrier therefore occurs at *r*_LxS_ ≈ 0 for all lipid species investigated, even though { *α*_*i*_} were determined specifically for DMPC. During all enhanced sampling simulations, *r*_LxS_ was calculated using a differentiable form for min(*d*_CC_),

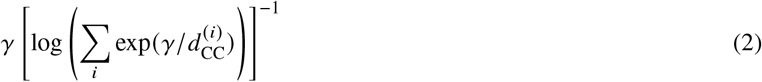

with *γ* = 200 nm, and for *n*_CC_,

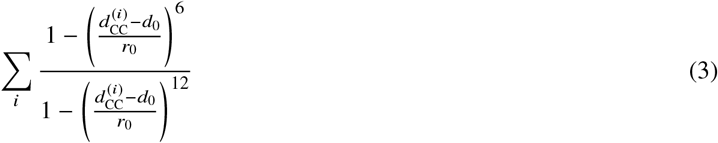

with *d*_0_ = 1 nm and *r*_0_ = 0.025 nm.

We performed umbrella sampling simulations (45) using the PLUMED 2 patch (46) for GROMACS to obtain the free energy profiles Δ *F* (*r*_LxS_) for each system. To generate initial configurations, a tagged lipid of the same species as the membrane lipids was randomly inserted into the solvent around each equilibrated bilayer such that the center of mass of the tagged lipid and of the bilayer were separated by at least 3.2 nm along *z*, which is the axis perpendicular to the bilayer. Each system was then energy minimized and equilibrated using the same two-step procedure as used for the bilayers with the addition of harmonic position restraints on the *z* coordinates of the tagged lipid’s heavy atoms with a force constant of 500 kJ/mol/nm^2^. Next, to generate initial configurations for each umbrella sampling window, a steered MD simulation was performed using a harmonic bias on *r*_LxS_ with a spring constant of 500 kJ/mol. Each umbrella sampling window was initialized with a configuration from the steered MD simulation that has a value of (*r*_LxS_) within 0.1 of the window’s bias center. Bias parameters used for umbrella sampling are tabulated in Table S1 of the Supplemental Information (SI). All windows were run for 52 ns. Convergence was assessed by monitoring how the numerical estimate of Δ *F* (*r*_LxS_) changed in time. For all systems, converged values of the free energy barriers for desorption and insertion were obtained after 20 ns (Fig. S1). Data from 20 − 52 ns of all windows was combined with the weighted histogram analysis method (WHAM) (47) to obtain Δ *F* (*r*_LxS_). Error bars were calculated as the standard error of Δ *F* (*r*_LxS_) estimated from 4 independent 8 ns blocks. As indicated in Fig. 1A, by setting Δ *F*(*r*_LxS_) to zero at the global free energy minimum, the barriers for lipid desorption and insertion are defined as

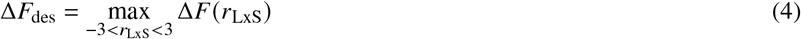

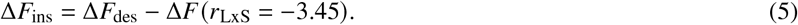

### Analysis of membrane properties

We analyzed trajectories of each bilayer system to assess how lipid chemistry and membrane phase influence the molecular structure of a membrane, and to provide a reference for rationalizing activation free energies for passive lipid transport in terms of physicochemical properties. In addition to the measures of hydrophobic lipid–membrane contacts defined above, we computed the membrane–solvent interaction energy, area per lipid, area of interfacial packing defects, membrane thickness, density profiles along the membrane normal, carbon–deuterium order parameters of the lipid tails, and radial distribution functions that characterize the intermolecular structure of the membrane–solvent interface. A combination of MDAnalysis (48) and NumPy (49) Python libraries in addition to GROMACS tools were used to calculate all properties. The membrane–solvent interaction energy, *E*_mem−solv_, was calculated as the sum of short-ranged Lennard-Jones and Coulomb interaction energy terms between the membrane and solvent. The average area per lipid, ⟨ *A*_lip_⟩, was calculated as the area of the box in the *xy* plane divided by the total number of lipids in a leaflet. Lipid packing defects, which are interfacial voids between polar lipid heads that expose aliphatic atoms to solvent, were identified with PackMem (50). The packing defect size constant, *π* _defect_, was obtained by fitting the distribution of defect areas, *A*_defect_, to a monoexponential decay,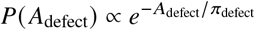. Fits were performed on *P*(*A*_defect_) ≥10^−4^ and *A*_defect_ ≥1.5 nm^2^ for L_*α*_ phase membranes or *A*_defect_ ≥0.5 nm^2^ for L_*β*_ phase membranes. Definitions of all other properties are provided in the SI. All analysis was performed on the final 300 ns of each bilayer trajectory split into 100 ns intervals for block averaging. Reported error is the standard error calculated from the three 100 ns intervals.

## RESULTS

For a series of 14 different single-component membranes, we quantified the free energy barriers that limit the rates of lipid desorption from and insertion into a membrane (Fig. 1). By calculating free energy profiles Δ*F* (*r*_LxS_) as a function of the reaction coordinate, which captures the collective motion of molecules that advances a transition (28, 51–53), we are able to extract dynamical information from them. Lipid desorption, which limits the rate of passive lipid transport, occurs when a lipid transitions from the membrane to the solvent, breaking hydrophobic contacts with the membrane along the way. Lipid insertion, which is the final step of lipid transfer, is the reverse process. The reaction coordinate for lipid (L) transport via solvent (xS), *r*_LxS_, measures the breakage and formation of hydrophobic lipid–membrane contacts through a linear combination of the minimum distance, min *d*_CC_, and number of close contacts, *n*_CC_, between hydrophobic carbons of the transferring lipid and membrane (Fig. 1B) (27). We assume that *r*_LxS_ serves as a good reaction coordinate for all lipids investigated as long as all hydrophobic carbons of each lipid species (Fig. 1C) are used to calculate *n*_CC_. *r*_LxS_ describes progress from configurations with the lipid in the membrane (which exhibit many hydrophobic lipid–membrane contacts and large positive values of *r*_LxS_) to configurations with the lipid in solution (which have very few if any hydrophobic lipid–membrane contacts and negative values of *r*_LxS_). Correspondingly, the rate of lipid desorption is limited by the free energy barrier Δ*F* _des_, which is the difference between Δ*F* (*r*_LxS_ ≈0) and the global free energy minimum found at large positive values of *r*_LxS_; the rate of lipid insertion is limited by the free energy barrier Δ*F* _ins_, which is the difference between Δ*F*(*r*_LxS_ ≈0) and the free energy at very negative values of *r*_LxS_ (Fig. 1A).

### Increasing lipid acyl chain length increases the desorption barrier

To investigate how the chemical structure of the lipid tails influences the rate of lipid transport, we simulated a series of L_*α*_ phase membranes composed of PC lipids with acyl chain lengths ranging from 14 to 18 carbons and degrees of unsaturation ranging from zero to two (Fig. 1C). This list included three fully saturated lipids that only differ in chain length (DMPC, DPPC, and DSPC); three lipids that only differ by degree of unsaturation (DOPC, SOPC, and DSPC); and a mixed-chain lipid (POPC).

The free energy profiles of each lipid exhibit the same qualitative features (Fig. 2A). This commonality is consistent with our assumption that PC lipids with different tails share the same biophysical mechanism of lipid transport, and that the generic transition pathway is well characterized by *r*_LxS_. Importantly, all of the free energy profiles exhibit a barrier at *r*_LxS_ ≈ 0. However, the free energy profiles are quantitatively different.

**Figure 2:**
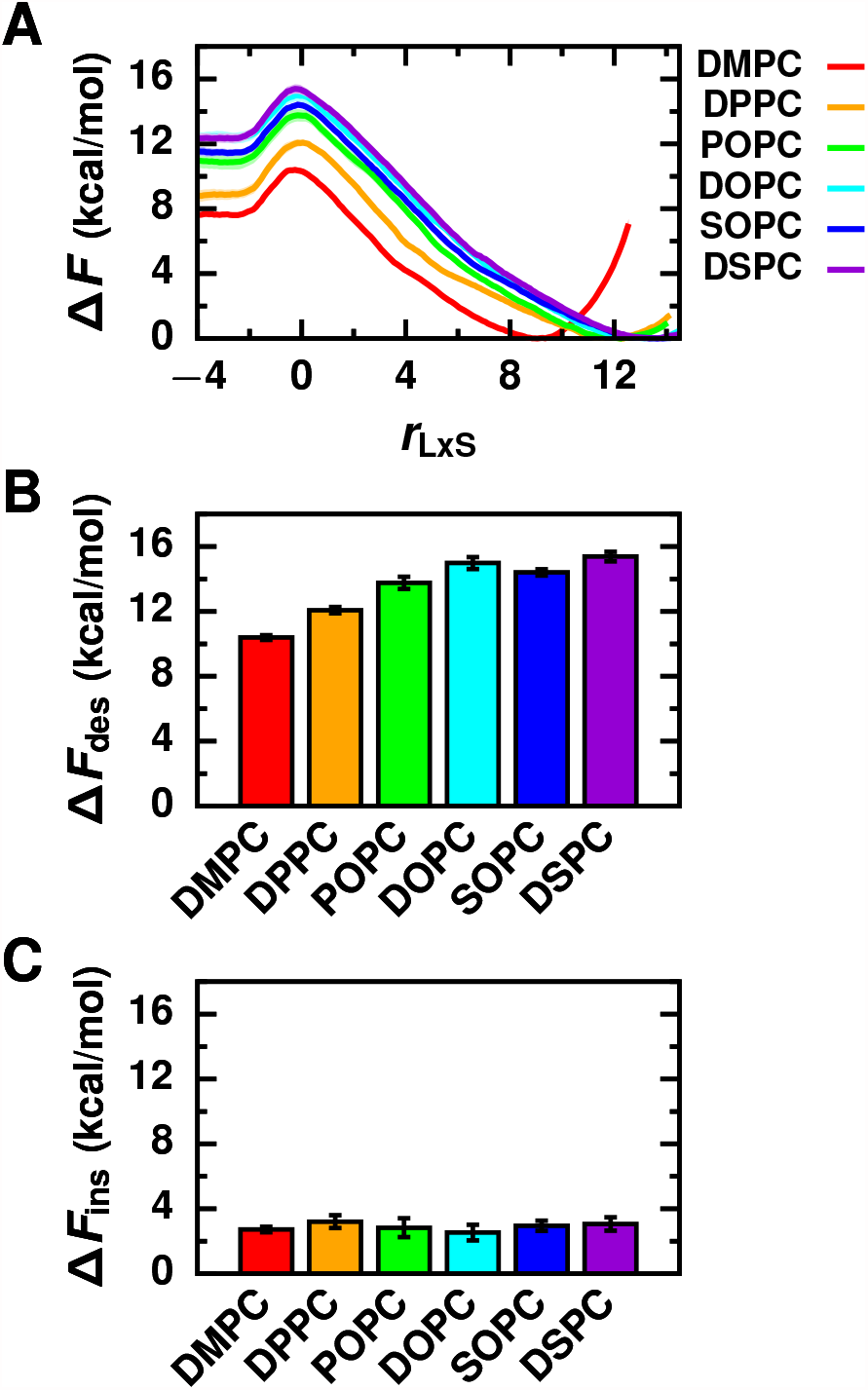
Increasing lipid acyl chain length increases the desorption barrier and does not substantially influence the insertion barrier. (A) Free energy profiles as a function of the reaction coordinate for L_*α*_ phase membranes composed of lipids with different tail chemistries. Corresponding free energy barriers are shown for (B) lipid desorption and (C) lipid insertion.

As shown in Fig. 2B, Δ*F* _des_ varies significantly with lipid tail chemistry, ranging from 10.4 ± 0.1 kcal/mol for DMPC to 15.4 ± 0.3 kcal/mol for DSPC (Table S2). Increasing the total number of carbons in the acyl chains increases Δ*F* _des_. Lipids with longer tails have more hydrophobic carbons (Fig. 1C) that can form hydrophobic contacts with surrounding lipids in the membrane (Table S3). Consequently, more hydrophobic lipid–membrane contacts must break for lipids with longer tails to desorb, explaining why (a) Δ*F* _des_ increases for the fully saturated lipids in the order DMPC < DPPC < DSPC, and (b) Δ*F* _des_ increases for the unsaturated lipids that differ in the length of a single acyl chain in the order POPC < SOPC. Δ*F* _des_ is smaller for lipids with unsaturated bonds than for fully saturated lipids with the same acyl chain lengths (Fig. 2B); both DOPC and SOPC have slightly reduced Δ*F* _des_ compared to DSPC. Membranes composed of lipids with increased degrees of unsaturation are more disordered (Fig. S2), with decreased bilayer thicknesses (Fig. S3 and Table S3) and increased areas per lipid (Table S3). As a result, hydrophobic contacts in these membranes can be more easily disrupted during lipid desorption.

Δ*F* _ins_ is substantially smaller than Δ*F* _des_, again demonstrating that lipid desorption limits the rate of lipid transport. In contrast to Δ*F* _des_, Δ*F* _ins_ does not vary significantly with lipid tail chemistry (Fig. 2C) and is 2.9 ± 0.4 kcal/mol on average (Table S2). During lipid insertion, a lipid must break through the membrane’s interface to form an initial hydrophobic lipid–membrane contact. Since all the lipids that we investigated with different tail chemistries have the same PC headgroup, the interfacial structure and chemistry of the membranes are quite similar (Fig. S3-S5). Consequently, the free energetic cost for a lipid to cross the membrane’s interface during insertion is roughly the same.

### Lipid headgroup influences both desorption and insertion barriers

To investigate how the chemical structure of the lipid headgroup influences lipid transport rates, we simulated a series of L_*α*_ phase membranes composed of PO lipids with both zwitterionic and anionic headgroups (Fig. 1D). This list included neutral lipids POPE, which has a terminal amine group, and POPC, which has a bulkier choline group; and the anionic lipids POPA and POPS, which have terminal acidic groups, and POPG and POPI, which have terminal polar groups.

The free energy profiles of the PO lipids with different headgroups exhibit the same general features as the profiles for the PC lipids, including a free energy barrier at *r*_LxS_ ≈ 0 (Fig. 3A). This commonality further supports our assumption that *r*_LxS_ is a generic reaction coordinate among glycerophospholipids. It additionally suggests that lipids with different headgroups and tails are transferred through the same biophysical mechanism, which is characterized by the breakage and formation of hydrophobic lipid–membrane contacts.

**Figure 3:**
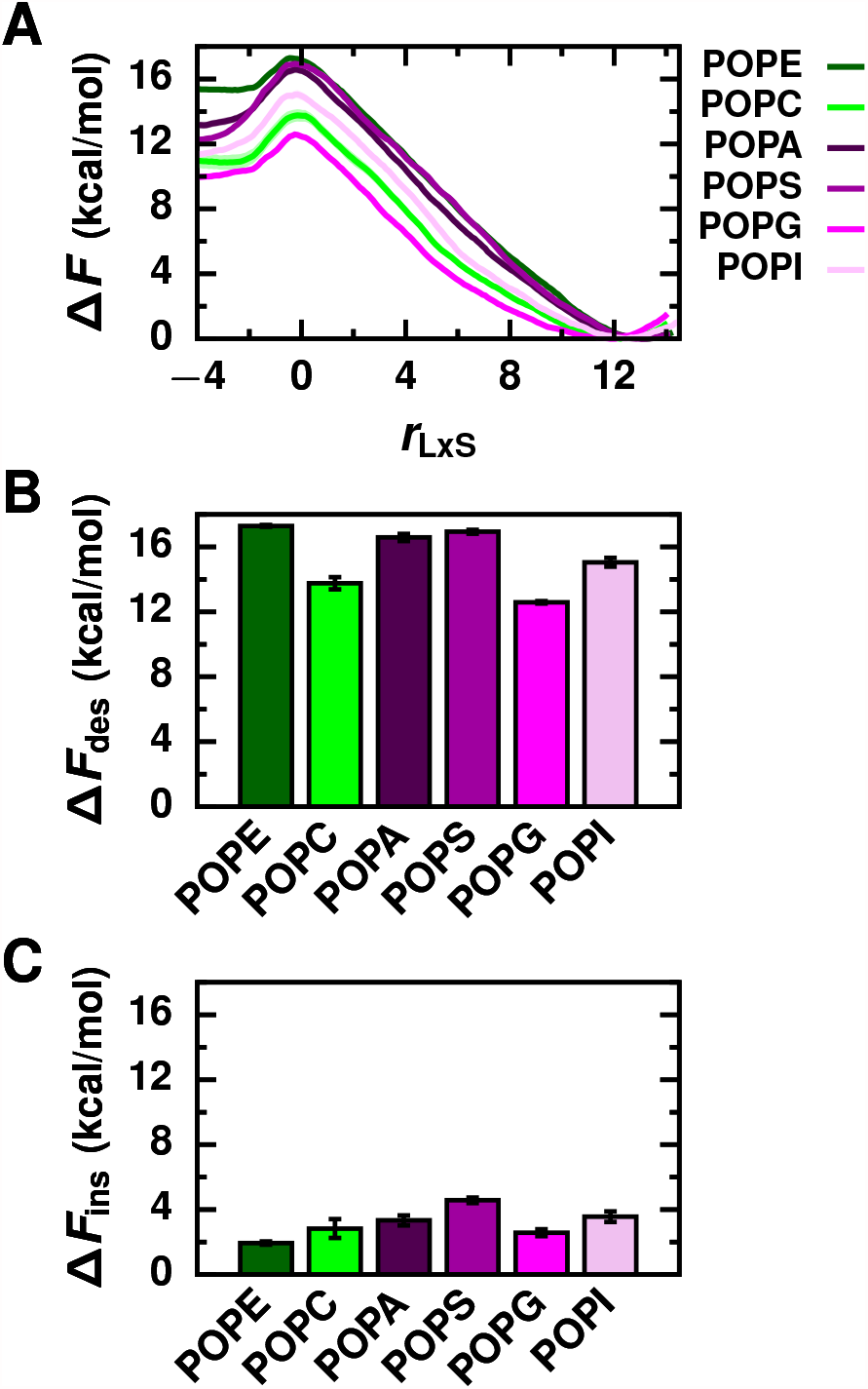
Lipid headgroup chemistry influences both desorption and insertion barriers. (A) Free energy profiles as a function of the reaction coordinate for L_*α*_ phase membranes composed of lipids with different headgroups. Corresponding free energy barriers are shown for (B) lipid desorption and (C) lipid insertion.

Quantitative differences in Δ*F* _des_ indicate that lipids with different headgroups are transported at different rates (Fig. 3B). Δ*F* _des_ ranges from 12.6 ± 0.1 kcal/mol for POPG to 17.3 ± 0.1 kcal/mol for POPE (Table S2). It is difficult to rationalize why certain headgroups result in smaller or larger Δ*F* _des_ based on their chemical structures alone. But, differences in Δ*F* _des_ can be explained based on each headgroup’s influence on general physical properties of a membrane (54–56), including lipid packing as measured by the average area per lipid (Table S3), membrane thickness (Fig. S3 and Table S3), and acyl chain order parameters (Fig. S2). Crucial to lipid transport, the headgroup impacts the number of hydrophobic contacts that a lipid forms with surrounding lipids in the membrane. A greater number of hydrophobic contacts between lipids in PE, PS, and PA membranes must be disrupted on average than in PI, PC, and PG membranes (Table S3) during lipid desorption. Δ*F* _des_ increases accordingly in the order POPG < POPC < POPI < POPA < POPS < POPE.

The interfacial structure and chemistry of a membrane is largely determined by the lipid headgroup (Fig. S3-S5). Δ*F* _ins_, which reflects the cost for a hydrophobic lipid tail to cross the membrane’s interface, thus varies with lipid headgroup (Fig. 3C), ranging from 1.9 ± 0.1 kcal/mol for POPE to 4.6 ± 0.2 kcal/mol for POPS (Table S2). On average, zwitterionic lipids have smaller Δ*F* _ins_ compared to anionic lipids; when comparing lipids with chemically similar headgroups (PE and PC; PA and PS; PG and PI), _ins_ decreases as the molecular size of the headgroup decreases. Decreasing the net charge of the membrane surface and decreasing the surface density of polar functional groups both reduce the free energetic cost of disrupting the membrane’s interfacial structure during the formation of an initial hydrophobic contact with an incoming lipid.

### Increasing membrane order increases both desorption and insertion barriers

Finally, we investigated how the rate of lipid transport depends on membrane phase. We simulated L_*β*_ phase membranes composed of the high melting temperature lipids DMPC, DPPC, and DSPC (Fig. 1E). Because the other lipid species investigated have very low melting temperatures, it would have been intractable to simulate them in a L_*β*_ phase and, furthermore, they are unlikely to be dominant components of highly ordered membrane domains at physiological temperatures.

Two different lipid arrangements are observed for the L_*β*_ phase membranes: Lipids in both leaflets of the DMPC and DPPC membranes tilt in the same direction (Fig. 1E), adopting an arrangement consistent with scattering experiments (57), whereas lipids in each leaflet of the DSPC membrane tilt in opposite directions (Fig. 1E), adopting a pleated arrangement that has been observed in other MD simulation studies (32, 58, 59). We do not expect the pleated arrangement of lipids in the L_*β*_ phase DSPC membrane to significantly impact the results reported below because other properties of the L_*β*_ phase DSPC membrane, such as the area per lipid and membrane thickness (Table S3), match experimental values (60–62), and because lipid desorption and insertion only involve one leaflet.

As with L_*α*_ phase membranes, the free energy profiles of L_*β*_ phase membranes exhibit a rate-limiting free energy barrier at *r*_LxS_ ≈ 0, albeit broader and more rugged (Fig. 4A). Thus, for both L_*α*_ and L_*β*_ phase membranes, the rate of lipid transport critically depends on the breakage and formation of hydrophobic lipid–membrane contacts. In contrast to free energy profiles of L_*α*_ phase membranes, profiles of L_*β*_ phase membranes exhibit a local free energy minimum at positive values of *r*_LxS_. At this local minimum, the transferring lipid adopts a splayed configuration with one tail anchored in the membrane and the other exposed to solvent (Fig. 4A and S6). While splayed intermediates have been observed in trajectories of lipid insertion into L_*α*_ phase membranes (27, 63), they are not locally thermodynamically stable in those cases. Given that splayed lipids form the transition state for vesicle fusion (64–69), the enhanced stability we have observed in more ordered membranes may help explain why ripple phase membranes fuse faster than L_*α*_ phase membranes in *in vitro* assays (70) and why many viral fusion proteins localize to lipid rafts (71–76). In L_*β*_ phase membranes, splayed lipids persist after hydrophobic contacts between a single lipid tail and the membrane are broken since a second free energy barrier must be crossed to break contacts with the other lipid tail. The free energy maximum at *r*_LxS_ ≈ 0 ultimately limits the rate of lipid transport; its value relative to the global minimum is reported as Δ*F*_des_ below.

**Figure 4:**
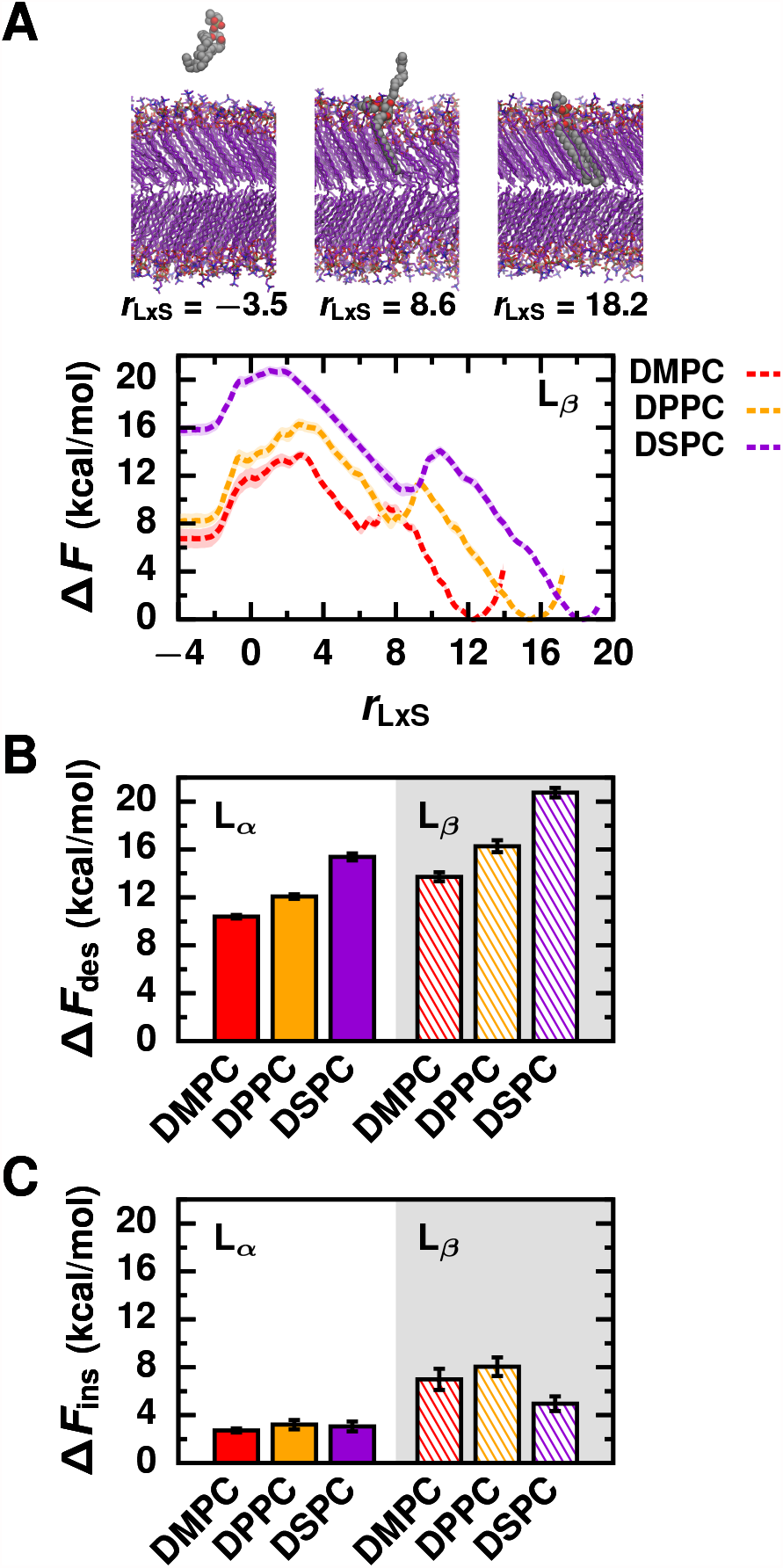
L_*β*_ phase membranes exhibit increased barriers for both desorption and insertion compared to L_*α*_ phase membranes. (A) Free energy profiles as a function of the reaction coordinate for L_*β*_ phase membranes, together with configurations of DSPC representating all three local minima of Δ*F*(*r*_LxS_): the lipid in solution, splayed lipid intermediate, and lipid in the membrane. (B and C) Free energy barriers of (B) lipid desorption and (C) lipid insertion for membranes simulated in both L_*α*_ and L_*β*_ phases.

Both Δ*F*_des_ and Δ*F*_ins_ are substantially larger for L_*β*_ compared to L_*α*_ phase membranes (Fig. 4B and 4C). For L_*β*_ membranes, Δ*F*_des_ ranges from 13.7 ± 0.4 kcal/mol for DMPC to 20.7 ± 0.4 kcal/mol for DSPC, and Δ*F*_ins_ ranges from 5.0 ± 0.6 kcal/mol for DSPC to 8.0 ± 0.8 kcal/mol for DPPC (Table S2). Lipids in highly ordered and tightly packed membranes, epitomized by L_*β*_ phase membranes, form a greater number of hydrophobic contacts with surrounding membrane lipids (Table S3). More contacts must be broken for a lipid to desorb from a L_*β*_ phase membrane compared to a L_*α*_ phase membrane, thus increasing Δ*F*_des_. Consistent with the trend observed for L_*α*_ phase membranes, L_*β*_ phase membranes composed of lipids with longer acyl chains also have larger Δ*F*_des_. The tight packing of lipids in L_*β*_ compared to L_*α*_ phase membranes also increases the density of polar lipid headgroups at the membrane’s interface (Fig. S3). Consequently, the free energetic cost for a lipid to traverse the membrane’s interface to form an initial hydrophobic lipid–membrane contact increases, explaining the overall increase in Δ*F*_ins_ for L_*β*_ phase membranes. Lipids pack more tightly in L_*β*_ phase DPPC and DMPC membranes than in L_*β*_ phase DSPC membranes, as indicated by their smaller areas (Table S3) and more ordered tails (Fig. S2). The resulting increased surface density of headgroups further hinders the disruption of the membrane’s interfacial structure during lipid insertion and, thus, Δ*F*_ins_ increases accordingly in the order DSPC < DMPC < DPPC for L_*β*_ phase membranes.

### Desorption barrier depends on a lipid’s local hydrophobic environment in a membrane

We attribute the main free energetic cost for lipid desorption to the disruption of a lipid’s locally hydrophobic environment in a membrane. A lipid’s hydrophobic environment is quantified by the average number of close hydrophobic contacts that a lipid in a membrane makes with surrounding lipids, ⟨*n*_CC_ ⟩_mem_. Because we consider any pair of hydrophobic carbons within 1 nm of each other as close contacts, lipids form hydrophobic contacts with both first and second nearest neighboring lipids (27). As shown in Fig. 5 and S7, a general trend between ⟨*n*_CC_⟩_mem_ and Δ*F*_des_ exists: As ⟨*n*_CC_⟩_mem_ increases, Δ*F*_des_ increases since more hydrophobic contacts must be broken to displace a lipid from the membrane. Δ*F*_des_ is roughly linearly correlated with ⟨*n*_CC_ ⟩_mem_, indicating that the cost of breaking each hydrophobic contact is approximately the same (roughly 5.8 cal/mol/contact). We note that Δ*F*_des_ is also roughly linearly correlated with the transfer free energy, Δ*F*_sol−mem_, which is the difference between Δ*F* (*r*_LxS_) at very negative values of *r*_LxS_ and the global free energy minimum of Δ*F* (*r*_LxS_) (Fig. S8). SinceΔ*F* _sol−mem_ characterizes the likelihood for a lipid to be fully solvated instead of fully within the membrane, Δ*F*_sol−mem_ is also highly indicative of membrane hydrophobicity. In contrast to Δ*F*_sol− mem_, ⟨*n*_CC_ ⟩_mem_ provides a molecular rationale for observed trends in Δ*F*_des_. Because ⟨*n*_CC_ ⟩_mem_ is sensitive to both lipid chemistry and membrane phase, the observations made above about how lipid tail chemistry, headgroup, and membrane phase individually affect Δ*F*_des_ are all attributable to differences in ⟨*n*_CC_ ⟩_mem_. In other words, ⟨*n*_CC_ ⟩_mem_ integrates the contributions to Δ*F*_des_ from all chemical and physical features of a membrane. Differences in Δ*F*_des_ between membranes that vary in both lipid chemistry and membrane phase, for example a L_*β*_ phase DMPC membrane compared to a L_*α*_ phase POPE membrane, are indeed explained by differences in ⟨*n*_CC_ ⟩_mem_. Thus, the hydrophobicity of a membrane’s core, as measured by ⟨*n*_CC_ ⟩_mem_, predominantly determines Δ*F*_des_ and, consequently, the rate of passive lipid transport.

**Figure 5:**
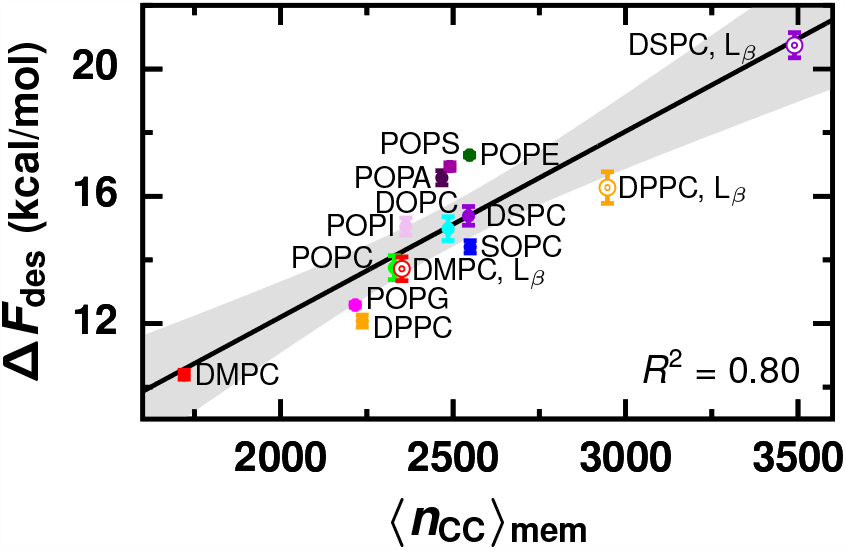
The barrier for desorption of a lipid is determined by its local hydrophobic membrane environment, which we quantify by the average number of close hydrophobic contacts between membrane lipids, ⟨*n*_CC_⟩_mem_. Black line is the best linear fit: Δ*F*_des_ = 0.0058⟨*n*_CC_⟩_mem_ + 0.53. Gray region indicates 95% confidence interval.

### Insertion barrier depends on a membrane’s interfacial hydrophobicity

We attribute the main free energetic cost for lipid insertion to the disruption of the membrane’s stratified chemical organization when the lipid breaches the membrane’s interface to form an initial hydrophobic lipid–membrane contact. The price of such a “mixing” of lipid headgroups and tails is greatest when the two groups are most chemically dissimilar; there is more resistance for lipid insertion into membranes with more hydrophilic surfaces. However, this picture is complicated by the fact that a membrane’s interface is chemically heterogeneous at the molecular scale, exhibiting regions dominated by headgroups and regions where tails are partially exposed to solvent. Contributions from both types of regions must be considered to explain all observed variations in Δ*F*_ins_. To compare different membranes, we characterize the net chemical character of their surfaces by an interfacial hydrophobicity. (The textbook description of lipid membranes’ surfaces is strongly hydrophilic, corresponding to very low interfacial hydrophobicity.) We quantify contributions from the headgroups by the average membrane–solvent interaction energy, ⟨*E*_mem−solv_⟩, and we consider membranes with less favorable ⟨*E*_mem−solv_⟩ to have more hydrophobic interfaces. As shown in Fig. 6A, for membranes with different headgroups, Δ*F*_ins_ decreases as ⟨*E*_mem−solv_⟩ becomes less favorable since the cost to compromise interactions between the headgroups and solvent during lipid insertion is reduced. For membranes with the same headgroup, differences in Δ*F*_ins_ arise from variations in the contributions to the interfacial hydrophobicity from regions where lipid tails are partially exposed. Because these regions result from lipid packing defects, we quantify these contributions by the relative area of packing defects, *Ā*_defect_, defined as the ratio of the characteristic packing defect size *π*_defect_ (50) (Fig. S9) to the average area of a lipid in the membrane, ⟨*A*_lip_ ⟩. We consider membranes with increased *Ā*_defect_ to have more hydrophobic interfaces. As shown in Fig. 6B, for membranes with the same headgroup but different tails or phases, Δ*F*_ins_ decreases as *Ā*_defect_ increases since the cost to locally expose the membrane’s hydrophobic core during lipid insertion is reduced.

**Figure 6:**
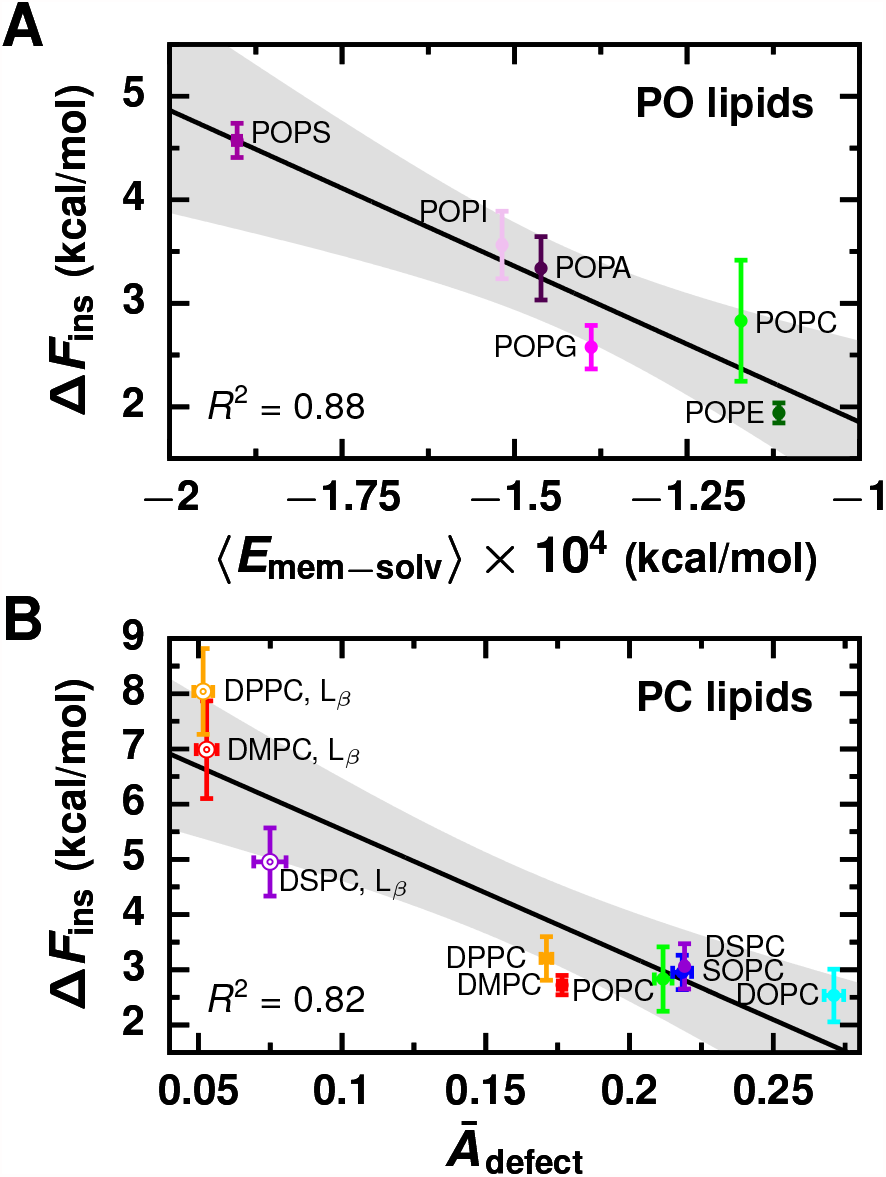
The barrier for lipid insertion is determined by a membrane’s interfacial hydrophobicity, which we quantify by the average membrane–solvent interaction energy (⟨*E*_mem− solv_ ⟩) and the relative area of membrane packing defects (*Ā*_defect_). (A) Changes in Δ*F*_ins_ due to lipid headgroup chemistry are attributed to differences in ⟨*E*_mem− solv_⟩. Black line is the best linear fit: Δ*F*_ins_ = −0.00030 ⟨*E*_mem− solv_⟩ −1.2. (B) Changes in Δ*F*_ins_ due to lipid tail chemistry and membrane phase are attributed to differences in *Ā*_defect_. Black line is the best linear fit: Δ*F*_ins_ = −22.9 *Ā*_defect_ + 7.8. Gray regions indicate 95% confidence intervals.

Overall, a membrane’s interfacial hydrophobicity determines Δ*F*_ins_, and, consequently, the rate of lipid insertion. To identify a relationship between Δ*F*_ins_ and quantitative metrics of a membrane’s interfacial hydrophobicity, we have considered variations due to lipid headgroup, which modulate interfacial hydrophobicity through changes in ⟨*E*_mem− solv_ ⟩, separately from variations due to tail chemistry and membrane phase, which tune interfacial hydrophobicity through changes in *Ā*_defect_. More sophisticated and precise measures of a membrane’s interfacial hydrophobicity, such as those developed for protein surfaces (77–80), may abrogate the need to consider contributions from headgroups separately from lipid tails and membrane phase. However, the two simple measures that we identify, ⟨*E*_mem− solv_ ⟩and *Ā*_defect_, reasonably explain variations in Δ*F*_ins_ for different membranes relative to the errors in calculated values of Δ*F*_ins_.

## DISCUSSION

### Passive lipid transport rate depends on membrane hydrophobicity

To decipher the physicochemical properties that control the rate of passive lipid transport between membranes, we systematically investigated how the free energy barriers for lipid desorption and insertion vary with lipid chemistry and membrane phase. To do so efficiently, we calculated free energy profiles as a function of the reaction coordinate *r*_LxS_, which we first identified for DMPC transport between L_*α*_ phase membranes (27) and here assumed applicable to similar systems. All of the free energy profiles exhibit a rate-limiting barrier at nearly the same value of the reaction coordinate (Fig. 2A, 3A, and 4A), supporting our assumption that *r*_LxS_ is a good reaction coordinate for all glycerophospholipids; regardless of glycerophospholipid chemistry or membrane phase, the dynamics of lipid transport are best described in terms of the breakage and formation of hydrophobic lipid–membrane contacts. In all cases, the free energy barrier for lipid desorption (Fig. 2B, 3B, and 4B) is at least twice the barrier for lipid insertion (Fig. 2C, 3C, and 4C), confirming that desorption limits the rate of lipid transport and that the free energy barrier for desorption is equivalent to the activation free energy barrier for lipid transport.

We have revealed a correlation between the activation free energy and the hydrophobicity of a lipid’s local membrane environment, as measured by the average number of close hydrophobic contacts that a lipid forms with surrounding membrane lipids, ⟨*n*_CC_ ⟩_mem_ (Fig. 5). ⟨*n*_CC_ ⟩_mem_, and hence the activation free energy, is sensitive to subtle changes in a membrane’s chemistry and structure. In general, lipids form more hydrophobic contacts in thick, tightly packed, and ordered membranes (Fig. S10). Membranes with these characteristics are typically composed of lipids with long, saturated acyl chains, lipids with headgroups that facilitate tight lipid packing, or lipids in ordered phases. Indeed, such membranes exhibit increased activation free energies (Fig. 2B, 3B, and 4B).

### Trends from simulation and experiment agree quantitatively

Our reported activation free energies significantly underestimate values estimated from experiments by at least 9 kcal/mol; of the membranes that have also been studied experimentally (L_*α*_ phase DMPC, DPPC, POPC, and DOPC; and L_*β*_ phase DMPC and DPPC), we calculated activation free energies ranging from 10.4 to 16.3 kcal/mol whereas experimental estimates range from 20.8 to 25.6 kcal/mol (13, 20, 23, 24, 26, 81) (Table S2). Due to the exponential dependence of the rate on the activation free energy, this discrepancy corresponds to a lipid transfer rate calculated from simulations that is roughly six-orders-of-magnitude faster than experimentally measured rates. As discussed in our previous study (27), this significant discrepancy may be due to any number of factors, including force field inaccuracies and differences in the solution ionic strength used in simulations and experiments. Despite this discrepancy in absolute activation free energies, differences in activation free energies due to specific changes in physicochemical properties calculated from our simulations agree quantitatively with experimental values.

### Influence of tail chemistry on the activation free energy

Lipids with long, saturated acyl chains exhibit increased activation free energies (Fig. 2B) and slower experimentally measured transfer rates (12–23, 25) than counterparts with unsaturated and shorter acyl chains. We report an average increase in the activation free energy of 0.62 kcal/mol per methylene unit added to the tails of a saturated lipid due to an average increase in ⟨*n*_CC_ ⟩_mem_ of 103 contacts per methylene unit. This corresponds to a roughly 10-fold decrease in the transfer rate per addition of two carbons to each acyl chain. Our results are in quantitative agreement with previous *in vitro* experiments that estimate an increase in the activation free energy of 0.32 −0.65 kcal/mol per methylene unit (15, 16, 19). Furthermore, our finding that the activation free energy depends on the hydrophobic environment around the desorbing lipid may explain why different values are reported from experiments using different donor membranes. Changing the desorbing lipid’s tail length alters the average number of hydrophobic contacts it makes with surrounding lipids, ⟨*n*_CC_ ⟩, and correspondingly alters its activation free energy. However, if donor membranes contain components capable of regulating ⟨*n*_CC_ ⟩_mem_, then desorbing lipids with different tail lengths may not exhibit substantially different ⟨*n*_CC_ ⟩, resulting in similar activation free energies. Proteins, for example, could act as such regulating components. Consistent with this hypothesis, smaller changes in the activation free energy were determined from experiments that utilized model high-density lipoproteins (HDL) composed of apolipoprotein A-I (apo A-I) and POPC as donors (16) compared to those using large unilamellar vesicles composed of only PC lipids (15, 19). Apo A-I wraps around lipids in model HDL to create a discoidal morphology (82), suggesting that it may regulate ⟨*n*_CC_ ⟩_mem_ by controlling the membrane’s overall structure. Alternatively, changes in ⟨*n*_CC_ ⟩_mem_ could be directly compensated by specific protein–lipid interactions.

### Influence of membrane phase on the activation free energy

Lipids in highly ordered L_*β*_ phase membranes exhibit the largest activation free energies (Fig. 4B) and slowest experimentally measured transfer rates (20, 22, 26). We report an average increase of 4.3 kcal/mol in the activation free energy for a L_*β*_ phase membrane compared to a L_*α*_ one due to an average increase in ⟨*n*_CC_ ⟩_mem_ of 762 contacts. This corresponds to a lipid transfer rate that is roughly three-orders-of-magnitude slower for L_*β*_ compared to L_*α*_ phase membranes. In agreement with our results, *in vitro* assays have demonstrated that lipid transport between L_*β*_ phase membranes has an increased activation free energy of 1.6 −12.7 kcal/mol compared to L_*α*_ phase membranes (20, 22, 26). We suggest that the wide range of activation free energies reported likely reflects differences in the hydrophobicity of the donor membranes used in the experiments. The smallest experimental difference is reported for lipid transport between highly curved small unilamellar vesicles (20), which are more disordered than planar membranes (83, 84). Thus, they likely have reduced ⟨*n*_CC_ ⟩_mem_, and correspondingly reduced activation free energy, compared to other experimental L_*β*_ phase systems (22, 26).

### Influence of headgroup on the activation free energy

We report an increase in the activation free energy for lipids with different headgroups in the order PG < PC < PA < PS < PE due to an increase in ⟨*n*_CC_ ⟩_mem_ in the same order. We cannot directly compare our results to experiments since significantly different donor membrane compositions were used. While we used donor membranes composed of the same lipid species as the one being transferred, donor membranes composed of predominantly PC lipids were used in experiments (12, 15, 17, 25). Nevertheless, we note that in multiple experiments, PE lipids are transferred at slower rates than PC lipids (15, 25), consistent with our results. However, no consensus ranking of transfer rates for lipids with other headgroups has been reached based on experiments (12, 15, 17, 25). Results from even the same experimental setup have often been difficult to interpret (15, 17, 25), most likely because no clear trend in the activation energy can be identified based on the chemical structure of the headgroup alone. Our finding that the activation free energy depends on ⟨*n*_CC_ ⟩_mem_, which depends on lipid headgroup chemistry in a nontrivial way, may help guide the design of experiments to conclusively assess the impact of headgroup on transfer rates.

### Influence of physicochemical properties on the free energy barrier for lipid insertion

In contrast to the activation free energy barrier for lipid desorption, trends in the free energy barrier for lipid insertion can be explained by chemical features of the headgroup, especially net charge, presence of terminal acidic groups, and molecular size (Fig. 3C). Changing any of these chemical features modulates the membrane’s interfacial hydrophobicity by altering the strength of membrane–solvent interactions (Table S3). A membrane’s interfacial hydrophobicity can also be tuned by changing lipid tail chemistry or membrane phase so that packing defects expose more of the hydrophobic core (Fig. S9 and Table S3). Membranes with increasingly hydrophobic surfaces, characterized by weaker headgroup–solvent interactions and larger packing defects, exhibit reduced insertion free energy barriers (Fig. 6) and correspondingly faster lipid insertion rates. Unfortunately, the rate of lipid insertion is too fast to monitor with standard experimental techniques, so we cannot compare our calculated free energy barriers for insertion to experimental ones. Nevertheless, one of the parameters that we find controls the rate of lipid insertion is similar to the membrane property that is predictive of experimentally measured rates of nanoparticle insertion into PC membranes: Nanoparticle insertion rates depend on the prevalence of low density areas at the membrane surface, which are physically similar to packing defects (85). Given this similarity, we suspect that accounting for changes in a membrane’s interfacial hydrophobicity through additional measures such as the membrane–solvent interaction energy may be necessary to predict nanoparticle insertion rates into more complex membranes that contain lipids with diverse headgroups and better mimic cell membrane compositions.

### Membrane hydrophobicity may help direct cellular lipid traffic

We investigated passive lipid transfer between single-component membranes; however, cellular membrane compositions are significantly more complex, containing not only a diversity of glycerophospholipids but also sterols, sphingolipids, and membrane proteins, for example (1, 2). We suspect that membrane hydrophobicity may determine the rate of passive lipid transport between both single- and multi-component membranes, although further work is required to test this hypothesis. Our observation that the rate depends on the physicochemical properties of the donor membrane within 1 nm of the transferring lipid suggests that transport rates can be regulated, for example, by local changes in membrane composition and membrane proteins that alter membrane hydrophobicity. Sequestering lipids into ordered, tightly packed lipid rafts (4), where a significant expenditure of energy is required to extract a lipid, may create pools of non-exchangeable lipids within a single membrane. Membrane physicochemical properties vary even more substantially between organelles, creating two cell membrane territories: Loose lipid packing and minimally charged membrane surfaces define the territory found among membranes of the early secretory pathway, whereas tight lipid packing and highly anionic membrane surfaces define the territory found among late secretory membranes (7, 86). Given that tight lipid packing increases the hydrophobicity of the membrane core and that stronger membrane electrostatics decrease interfacial hydrophobicity, we suggest that the free energy barriers for lipid transport may transition from low to high between these two territories such that lipids would passively transfer among early secretory membranes faster than they would among late secretory membranes.

Within cells, lipid transfer proteins are largely responsible for specifically transporting individual lipids between membranes (9, 10, 87). Our findings about passive lipid transport suggest that lipid transfer proteins may generally enhance the rate of lipid desorption by disrupting the hydrophobic environment around a target lipid. Those that transport lipids from membranes with highly hydrophobic cores, such as may be found among late secretory membranes, may have acquired additional mechanisms to overcome increased activation free energies. Similarly, lipid transfer proteins may aid lipid insertion by increasing the surface hydrophobicity of the membrane. For example, multiple lipid transfer proteins contain basic surface regions that enhance binding to anionic membranes (88–90) and that could also compensate for the disruption of especially favorable interactions between solvent and anionic lipids during insertion. Thus, the membrane physicochemical properties that control the rate of passive lipid transport may also be exploited to direct non-vesicular lipid traffic *via* lipid transfer proteins.

## AUTHOR CONTRIBUTIONS

J.R.R. designed the research. J.R.R. and G.E.G. carried out all simulations and analyzed the data, with guidance from P.L.G.

J.R.R. and P.L.G. wrote the article.

## ACKNOWLEDGMENTS

J.R.R. acknowledges the support of the National Science Foundation Graduate Research Fellowship Program under Grant No. DGE 1752814. G.E.G. was supported by the UC Berkeley College of Chemistry’s Summer Undergraduate Research Program. P.L.G. was supported by the Director, Office of Basic Energy Sciences, Office of Science, US Department of Energy under Contract DE-AC02-05CH11231, through the Chemical Sciences Division of Lawrence Berkeley National Laboratory. This work used computational resources from the Extreme Science and Engineering Discovery Environment (XSEDE), which is supported by National Science Foundation grant number ACI-1548562, and from Berkeley Laboratory Research Computing.

## SUPPORTING CITATIONS

Reference (91) appears in the Supplemental Information.

## SUPPLEMENTAL INFORMATION

An online supplement to this article can be found by visiting BJ Online at http://www.biophysj.org.

## REFERENCES

1. Harayama, T., and H. Riezman, 2018. Understanding the diversity of membrane lipid composition. Nat. Rev. Mol. Cell Biol. 19:281–296.

2. van Meer, G., D. R. Voelker, and G. W. Feigenson, 2008. Membrane lipids: Where they are and how they behave. Nat. Rev. Mol. Cell Biol. 9:112–124.

3. Lorent, J. H., K. R. Levental, L. Ganesan, G. Rivera-Longsworth, E. Sezgin, M. Doktorova, E. Lyman, and I. Levental, 2020. Plasma membranes are asymmetric in lipid unsaturation, packing and protein shape. Nat. Chem. Biol. 16:644–652.

4. Sezgin, E., I. Levental, S. Mayor, and C. Eggeling, 2017. The mystery of membrane organization: Composition, regulation and roles of lipid rafts. Nat. Rev. Mol. Cell Biol. 18:361–374.

5. de Mendoza, D., and M. Pilon, 2019. Control of membrane lipid homeostasis by lipid-bilayer associated sensors: A mechanism conserved from bacteria to humans. Prog. Lipid Res. 76:100996.

6. Ernst, R., S. Ballweg, and I. Levental, 2018. Cellular mechanisms of physicochemical membrane homeostasis. Curr. Opin. Cell Biol. 53:44–51.

7. Holthuis, J. C. M., and A. K. Menon, 2014. Lipid landscapes and pipelines in membrane homeostasis. Nature 510:48–57.

8. Jackson, C. L., L. Walch, and J. M. Verbavatz, 2016. Lipids and their trafficking: An integral part of cellular organization. Dev. Cell 39:139–153.

9. Lev, S., 2010. Non-vesicular lipid transport by lipid-transfer proteins and beyond. Nat. Rev. Mol. Cell Biol. 11:739–750.

10. Wong, L. H., A. T. Gatta, and T. P. Levine, 2019. Lipid transfer proteins: The lipid commute via shuttles, bridges and tubes. Nat. Rev. Mol. Cell Biol. 20:85–101.

11. Chiapparino, A., K. Maeda, D. Turei, J. Saez-Rodriguez, and A.-C. Gavin, 2016. The orchestra of lipid-transfer proteins at the crossroads between metabolism and signaling. Prog. Lipid Res. 61:30–39.

12. De Cuyper, M., and M. Joniau, 1985. Spontaneous intervesicular transfer of anionic phospholipids differing in the nature of their polar headgroup. Biochim. Biophys. Acta 814:374–380.

13. Pownall, H. J., D. L. Hickson, and L. C. Smith, 1983. Transport of biological lipophiles: Effect of lipophile structure. J. Am. Chem. Soc. 105:2440–2445.

14. Ferrell, J. E., K. J. Lee, and W. H. Huestis, 1985. Lipid transfer between phosphatidylcholine vesicles and human erythrocytes: Exponential decrease in rate with increasing acyl chain length. Biochemistry 24:2857–2864.

15. Homan, R., and H. J. Pownall, 1988. Transbilayer diffusion of phospholipids: Dependence on headgroup structure and acyl chain-length. Biochim. Biophys. Acta 938:155–166.

16. Pownall, H. J., D. L. M. Bick, and J. B. Massey, 1991. Spontaneous phospholipid transfer: Development of a quantitative model. Biochemistry 30:5696–5700.

17. Richens, J. L., A. I. I. Tyler, H. M. G. Barriga, J. P. Bramble, R. V. Law, N. J. Brooks, J. M. Seddon, O. Ces, and P. O’Shea, 2017. Spontaneous charged lipid transfer between lipid vesicles. Sci. Rep. 7:12606.

18. Nichols, J. W., 1985. Thermodynamics and kinetics of phospholipid monomer vesicle interaction. Biochemistry 24:6390–6398.

19. Silvius, J. R., and R. Leventis, 1993. Spontaneous interbilayer transfer of phospholipids: Dependence of acyl chain composition. Biochemistry 32:13318–13326.

20. McLean, L. R., and M. C. Phillips, 1984. Kinetics of phosphatidylcholine and lysophosphatidylcholine exchange between unilamellar vesicles. Biochemistry 23:4624–4630.

21. Bayerl, T. M., C. F. Schmidt, and E. Sackmann, 1988. Kinetics of symmetric and asymmetric phospholipid transfer between small sonicated vesicles studied by high-sensivity differential scanning calorimetry, NMR, electron-microscopy, and dynamic light-scattering. Biochemistry 27:6078–6085.

22. Massey, J. B., A. M. Gotto, and H. J. Pownall, 1982. Kinetics and mechanism of the spontaneous transfer of fluorescent phosphatidylcholines between apolipoprotein-phospholipid recombinants. Biochemistry 21:3630–3636.

23. Xia, Y., M. Li, K. Charubin, Y. Liu, F. A. Heberle, J. Katsaras, B. X. Jing, Y. X. Zhu, and M. P. Nieh, 2015. Effects of nanoparticle morphology and acyl chain length on spontaneous lipid transfer rates. Langmuir 31:12920–12928.

24. Jones, J. D., and T. E. Thompson, 1990. Mechanism of spontaneous, concentration-dependent phospholipid transfer between bilayers. Biochemistry 29:1593–1600.

25. Nichols, J. W., and R. E. Pagano, 1982. Use of resonance energy-transfer to study the kinetics of amphiphile transfer between vesicles. Biochemistry 21:1720–1726.

26. Wimley, W. C., and T. E. Thompson, 1990. Exchange and flip-flop of dimyristoylphosphatidylcholine in liquid-crystalline, gel, and 2-component, 2-phase large unilamellar vesicles. Biochemistry 29:1296–1303.

27. Rogers, J. R., and P. L. Geissler, 2020. Breakage of hydrophobic contacts limits the rate of passive lipid exchange between membranes. J. Phys. Chem. B 124:5884–5898.

28. Zuckerman, D. M., 2010. Statistical physics of biomolecules: An introduction. CRC Press, Boca Raton, FL.

29. Jo, S., T. Kim, V. G. Iyer, and W. Im, 2008. CHARMM-GUI: A web-based graphical user interface for CHARMM. J. Comput. Chem. 29:1859–1865.

30. Wu, E. L., X. Cheng, S. Jo, H. Rui, K. C. Song, E.M. Dávila-Contreras, Y. Qi, J. Lee, V. Monje-Galvan, R. M. Venable, J. B. Klauda, and W. Im, 2014. CHARMM-GUI membrane builder toward realistic biological membrane simulations. J. Comput. Chem. 35:1997–2004.

31. Nitschke, N., K. Atkovska, and J. S. Hub, 2016. Accelerating potential of mean force calculations for lipid membrane permeation: System size, reaction coordinate, solute-solute distance, and cutoffs. J. Chem. Phys. 145:125101.

32. Khakbaz, P., and J. B. Klauda, 2018. Investigation of phase transitions of saturated phosphocholine lipid bilayers via molecular dynamics simulations. Biochm. Biophys. Acta Biomembr. 1860:1489–1501.

33. Domański, J., P. J. Stansfeld, M. S. P. Sansom, and O. Beckstein, 2010. Lipidbook: A public repository for force-field parameters used in membrane simulations. J. Membr. Biol. 236:255–258.

34. Klauda, J. B., R. M. Venable, J. A. Freites, J. W. O’Connor, D. J. Tobias, C. Mondragon-Ramirez, I. Vorobyov, A. D. MacKerell, and R. W. Pastor, 2010. Update of the CHARMM all-atom additive force field for lipids: Validation on six lipid types. J. Phys. Chem. B 114:7830–7843.

35. MacKerell, A. D., D. Bashford, M. Bellott, R. L. Dunbrack, J. D. Evanseck, M. J. Field, S. Fischer, J. Gao, H. Guo, S. Ha, D. Joseph-McCarthy, L. Kuchnir, K. Kuczera, F. T. K. Lau, C. Mattos, S. Michnick, T. Ngo, D. T. Nguyen, B. Prodhom, W. E. Reiher, B. Roux, M. Schlenkrich, J. C. Smith, R. Stote, J. Straub, M. Watanabe, J. Wiórkiewicz-Kuczera, D. Yin, and M. Karplus, 1998. All-atom empirical potential for molecular modeling and dynamics studies of proteins. J. Phys. Chem. B 102:3586–3616.

36. Abraham, M. J., T. Murtola, R. Schulz, S. Páll, J. C. Smith, B. Hess, and E. Lindahl, 2015. GROMACS: High performance molecular simulations through multi-level parallelism from laptops to supercomputers. SoftwareX 1-2:19–25.

37. Nosé, S., 1984. A molecular-dynamics method for simulations in the canonical ensemble. Mol. Phys. 52:255–268.

38. Hoover, W. G., 1985. Canonical dynamics - equilibrium phase-space distributions. Phys. Rev. A 31:1695–1697.

39. Hockney, W. R., 1970. The potential calculation and some applications. Methods Comput. Phys. 9:136–211.

40. Hess, B., H. Bekker, H. J. Berendsen, and J. G. Fraaije, 1997. Lincs: A linear constraint solver for molecular simulations. J. Comput. Chem. 18:1463–1472.

41. Essmann, U., L. Perera, M. L. Berkowitz, T. Darden, H. Lee, and L. G. Pedersen, 1995. A smooth particle mesh ewald method. J. Chem. Phys. 103:8577–8593.

42. Páll, S., and B. Hess, 2013. A flexible algorithm for calculating pair interactions on simd architectures. Comput. Phys. Commun. 184:2641–2650.

43. Berendsen, H. J. C., J. P. M. Postma, W. F. van Gunsteren, A. DiNola, and J. R. Haak, 1984. Molecular dynamics with coupling to an external bath. J. Chem. Phys. 81:3684–3690.

44. Parrinello, M., and A. Rahman, 1981. Polymorphic transitions in single-crystals - a new molecular-dynamics method. J. Appl. Phys. 52:7182–7190.

45. Torrie, G. M., and J. P. Valleau, 1977. Non-physical sampling distributions in monte-carlo free-energy estimation - umbrella sampling. J. Comput. Phys. 23:187–199.

46. Tribello, G. A., M. Bonomi, D. Branduardi, C. Camilloni, and G. Bussi, 2014. PLUMED 2: New feathers for an old bird. Comput. Phys. Commun. 185:604–613.

47. Kumar, S., D. Bouzida, R. H. Swendsen, P. A. Kollman, and J. M. Rosenberg, 1992. The weighted histogram analysis method for free-energy calculations on biomolecules. 1. The method. J. Comput. Chem. 13:1011–1021.

48. Michaud-Agrawal, N., E. J. Denning, T. B. Woolf, and O. Beckstein, 2011. MDAnalysis: A toolkit for the analysis of molecular dynamics simulations. J. Comput. Chem. 32:2319–2327.

49. Harris, C. R., K. J. Millman, S. J. van der Walt, R. Gommers, P. Virtanen, D. Cournapeau, E. Wieser, J. Taylor, S. Berg, N. J. Smith, R. Kern, M. Picus, S. Hoyer, M. H. van Kerkwijk, M. Brett, A. Haldane, J. F. del Río, M. Wiebe, P. Peterson, P. Gérard-Marchant, K. Sheppard, T. Reddy, W. Weckesser, H. Abbasi, C. Gohlke, and T. E. Oliphant, 2020. Array programming with numpy. Nature 585:357–362.

50. Gautier, R., A. Bacle, M. L. Tiberti, P. F. Fuchs, S. Vanni, and B. Antonny, 2018. Packmem: A versatile tool to compute and visualize interfacial packing defects in lipid bilayers. Biophys. J. 115:436–444.

51. Du, R., V. S. Pande, A. Y. Grosberg, T. Tanaka, and E. S. Shakhnovich, 1998. On the transition coordinate for protein folding. J. Chem. Phys. 108:334–350.

52. Dellago, C., P. G. Bolhuis, and P. L. Geissler, 2002. Transition path sampling. In I. Prigogine, and S. A. Rice, editors, Advances in chemical physics, John Wiley & Sons, New York, volume 123, 1–78.

53. Bolhuis, P. G., D. Chandler, C. Dellago, and P. L. Geissler, 2002. Transition path sampling: Throwing ropes over rough mountain passes, in the dark. Annu. Rev. Phys. Chem. 53:291–318.

54. Seu, K. J., L. R. Cambrea, R. M. Everly, and J. S. Hovis, 2006. Influence of lipid chemistry on membrane fluidity: Tail and headgroup interactions. Biophys. J. 91:3727–3735.

55. Dickey, A., and R. Faller, 2008. Examining the contributions of lipid shape and headgroup charge on bilayer behavior. Biophys. J. 95:2636–2646.

56. Vamparys, L., R. Gautier, S. Vanni, W. F. D. Bennett, D. P. Tieleman, B. Antonny, C. Etchebest, and P. F. J. Fuchs, 2013. Conical lipids in flat bilayers induce packing defects similar to that induced by positive curvature. Biophys. J. 104:585–593.

57. Marsh, D., 2012. Lateral order in gel, subgel and crystalline phases of lipid membranes: Wide-angle x-ray scattering. Chem. Phys. Lipids 165:59–76.

58. Leekumjorn, S., and A. K. Sum, 2007. Molecular studies of the gel to liquid-crystalline phase transition for fully hydrated DPPC and DPPE bilayers. Biochim. Biophys. Acta 1768:354–365.

59. Tu, K., D. J. Tobias, J. K. Blasie, and M. L. Klein, 1996. Molecular dynamics investigation of the structure of a fully hydrated gel-phase dipalmitoylphosphatidylcholine bilayer. Biophys. J. 70:595–608.

60. Sun, W. J., S. Tristram-Nagle, R. M. Suter, and J. F. Nagle, 1996. Structure of gel phase saturated lecithin bilayers: Temperature and chain length dependence. Biophys. J. 71:885–891.

61. Tardieu, A., V. Luzzati, and F. C. Reman, 1973. Structure and polymorphism of the hydrocarbon chains of lipids: A study of lecithin-water phases. J. Mol. Biol. 75:711–733.

62. Lis, L. J., M. McAlister, N. Fuller, R. P. Rand, and V. A. Parsegian, 1982. Interactions between neutral phospholipid bilayer membranes. Biophys. J. 37:657–665.

63. Vermaas, J. V., and E. Tajkhorshid, 2014. A microscopic view of phospholipid insertion into biological membranes. J. Phys. Chem. B 118:1754–1764.

64. Kasson, P. M., E. Lindahl, and V. S. Pande, 2010. Atomic-resolution simulations predict a transition state for vesicle fusion defined by contact of a few lipid tails. PLoS Comput. Biol. 6:e1000829.

65. Smirnova, Y. G., S. J. Marrink, R. Lipowsky, and V. Knecht, 2010. Solvent-exposed tails as prestalk transition states for membrane fusion at low hydration. J. Am. Chem. Soc. 132:6710–6718.

66. Mirjanian, D., A. N. Dickey, J. H. Hoh, T. B. Woolf, and M. J. Stevens, 2010. Splaying of aliphatic tails plays a central role in barrier crossing during liposome fusion. J. Phys. Chem. B 114:11061–11068.

67. Stevens, M. J., J. H. Hoh, and T. B. Woolf, 2003. Insights into the molecular mechanism of membrane fusion from simulation: Evidence for the association of splayed tails. Phys. Rev. Lett. 91:188102.

68. Ohta-Iino, S., M. Pasenkiewicz-Gierula, Y. Takaoka, H. Miyagawa, K. Kitamura, and A. Kusumi, 2001. Fast lipid disorientation at the onset of membrane fusion revealed by molecular dynamics simulations. Biophys. J. 81:217–224.

69. Holopainen, J. M., J. Y. Lehtonen, and P. K. Kinnunen, 1999. Evidence for the extended phospholipid conformation in membrane fusion and hemifusion. Biophys. J. 76:2111–2120.

70. Lee, D. E., M. G. Lew, and D. J. Woodbury, 2013. Vesicle fusion to planar membranes is enhanced by cholesterol and low temperature. Chem. Phys. Lipids 166:45–54.

71. Liao, Z., L. M. Cimakasky, R. Hampton, D. H. Nguyen, and J. E. K. Hildreth, 2001. Lipid rafts and HIV pathogenesis: Host membrane cholesterol is required for infection by HIV type 1. AIDS Res. Hum. Retrovir. 17:1009–1019.

72. Klug, Y. A., E. Rotem, R. Schwarzer, and Y. Shai, 2017. Mapping out the intricate relationship of the HIV envelope protein and the membrane environment. Biochm. Biophys. Acta Biomembr. 1859:550–560.

73. Takeda, M., G. P. Leser, C. J. Russell, and R. A. Lamb, 2003. Influenza virus hemagglutinin concentrates in lipid raft microdomains for efficient viral fusion. Proc. Natl. Acad. Sci. U. S. A. 100:14610–14617.

74. Bavari, S., C. M. Bosio, E. Wiegand, G. Ruthel, A. B. Will, T. W. Geisbert, M. Hevey, C. Schmaljohn, A. Schmaljohn, and M. J. Aman, 2002. Lipid raft microdomains: A gateway for compartmentalized trafficking of Ebola and Marburg viruses. J. Exp. Med. 195:593–602.

75. Freitas, M. S., C. Follmer, L. T. Costa, C. Vilani, M. L. Bianconi, C. A. Achete, and J. L. Silva, 2011. Measuring the strength of interaction between the Ebola fusion peptide and lipid rafts: Implications for membrane fusion and virus infection. PLoS One 6:e15756–e15756.

76. Fecchi, K., S. Anticoli, D. Peruzzu, E. Iessi, M. C. Gagliardi, P. Matarrese, and A. Ruggieri, 2020. Coronavirus interplay with lipid rafts and autophagy unveils promising therapeutic targets. Front. Microbiol. 11:1821.

77. Patel, A. J., and S. Garde, 2014. Efficient method to characterize the context-dependent hydrophobicity of proteins. J. Phys. Chem. B 118:1564–1573.

78. Patel, A. J., P. Varilly, S. N. Jamadagni, M. F. Hagan, D. Chandler, and S. Garde, 2012. Sitting at the edge: How biomolecules use hydrophobicity to tune their interactions and function. J. Phys. Chem. B 116:2498–2503.

79. Rego, N. B., E. Xi, and A. J. Patel, 2019. Protein hydration waters are susceptible to unfavorable perturbations. J. Am. Chem. Soc. 141:2080–2086.

80. Rego, N. B., E. Xi, and A. J. Patel, 2021. Identifying hydrophobic protein patches to inform protein interaction interfaces. Proc. Natl. Acad. Sci. U. S. A. 118:e2018234118.

81. Nakano, M., M. Fukuda, T. Kudo, H. Endo, and T. Handa, 2007. Determination of interbilayer and transbilayer lipid transfers by time-resolved small-angle neutron scattering. Phys. Rev. Lett. 98:238101.

82. Brouillette, C. G., G. M. Anantharamaiah, J. A. Engler, and D. W. Borhani, 2001. Structural models of human apolipoprotein A-I: A critical analysis and review. Biochim. Biophys. Acta Mol. Cell Biol. Lipids 1531:4–46.

83. Lentz, B. R., Y. Barenholz, and T. E. Thompson, 1976. Fluorescence depolarization studies of phase transitions and fluidity in phospholipid bilayers. 1. Single component phosphatidylcholine liposomes. Biochemistry 15:4521–4528.

84. Yesylevskyy, S. O., T. Rivel, and C. Ramseyer, 2017. The influence of curvature on the properties of the plasma membrane. Insights from atomistic molecular dynamics simulations. Sci. Rep. 7:16078.

85. Liu, C., P. Elvati, S. Majumder, Y. Wang, A. P. Liu, and A. Violi, 2019. Predicting the time of entry of nanoparticles in lipid membranes. ACS Nano 13:10221–10232.

86. Bigay, J., and B. Antonny, 2012. Curvature, lipid packing, and electrostatics of membrane organelles: Defining cellular territories in determining specificity. Dev. Cell 23:886–895.

87. Wong, L. H., A. Copic, and T. P. Levine, 2017. Advances on the transfer of lipids by lipid transfer proteins. Trends Biochem. Sci. 42:516–530.

88. Iaea, D. B., I. Dikiy, I. Kiburu, D. Eliezer, and F. R. Maxfield, 2015. STARD4 membrane interactions and sterol binding. Biochemistry 54:4623–4636.

89. Watanabe, Y., Y. Tamura, S. Kawano, and T. Endo, 2015. Structural and mechanistic insights into phospholipid transfer by Ups1–Mdm35 in mitochondria. Nat. Commun. 6:7922.

90. Rogaski, B., and J. B. Klauda, 2012. Membrane-binding mechanism of a peripheral membrane protein through microsecond molecular dynamics simulations. J. Mol. Biol. 423:847–861.

91. Pluhackova, K., S. A. Kirsch, J. Han, L. P. Sun, Z. Y. Jiang, T. Unruh, and R. A. Bockmann, 2016. A critical comparison of biomembrane force fields: Structure and dynamics of model DMPC, POPC, and POPE bilayers. J. Phys. Chem. B 120:3888–3903.

