## Supplemental Information for "Membrane Hydrophobicity Determines the Activation Free Energy of Passive Lipid Transport"

### SUPPLEMENTAL METHODS

#### Analysis of membrane properties

*Membrane thickness.* The average membrane thickness was calculated as the average distance in $z$ between the phosphorous atoms of the top and bottom leaflets.

*Density profiles.* The density of various groups along the membrane normal relative to the center of mass of the membrane at $z = 0$ were calculated with the GROMACS tool `gmx density`. The groups are: (1) polar, which includes all atoms present in the system that are not hydrophobic carbons; (2) solvent, which includes all water molecules and any sodium ions present; (3) membrane, which includes all lipid atoms; (4) polar membrane, which includes all lipid atoms that are not hydrophobic carbons; and (5) hydrophobic, which includes all hydrophobic carbons.

*Acyl chain order parameters.* Deuterium order parameters for carbons in the lipid tails, except the ester and terminal carbons, were calculated with the GROMACS tool `gmx order`. The order parameters of carbons in double bonds were calculated following the instructions provided by Pluhackova et al. (1).

*Radial distribution functions.* To characterize the intermolecular structure of the membrane–solvent interface, we calculated radial distribution functions between the oxygen atoms of water molecules and either the phosphorous atoms or carbonyl oxygen atoms of the lipids using the GROMACS tool `gmx rdf`. Additionally, 2D radial distribution functions in the $xy$-plane were calculated between the phosphorous atoms of lipids in each leaflet individually and then averaged to obtain one 2D radial distribution function for each membrane.

#### SUPPLEMENTAL FIGURES

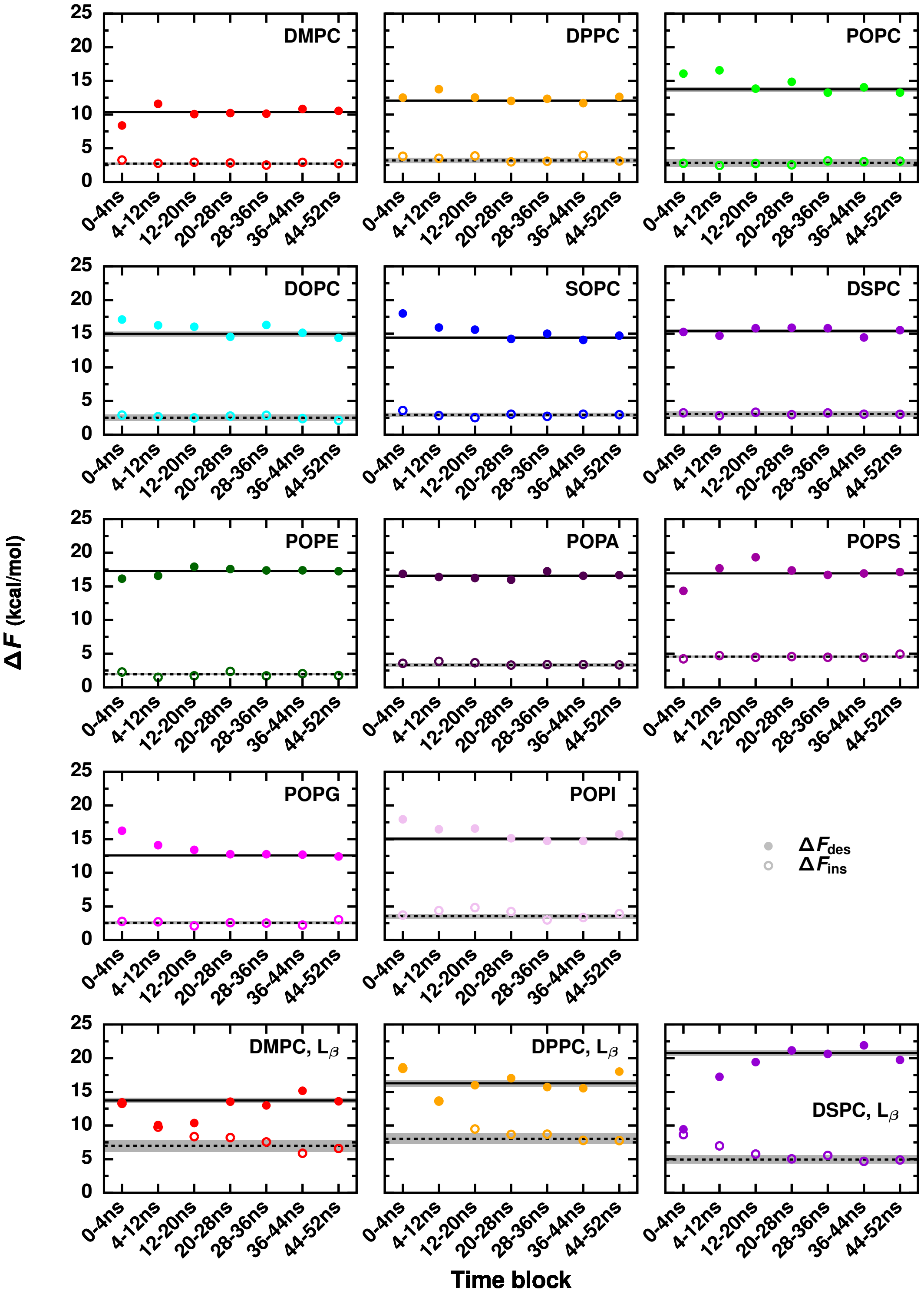


**Figure S1. Convergence of free energy calculations.** $\Delta F_{\mathrm{des}}$ and $\Delta F_{\mathrm{ins}}$ estimated from different simulation time blocks for each membrane. Final reported values of $\Delta F_{\mathrm{des}}$ and $\Delta F_{\mathrm{ins}}$ estimated from $20-52$ ns are plotted in black solid and dashed lines, respectively, with error bars in gray.

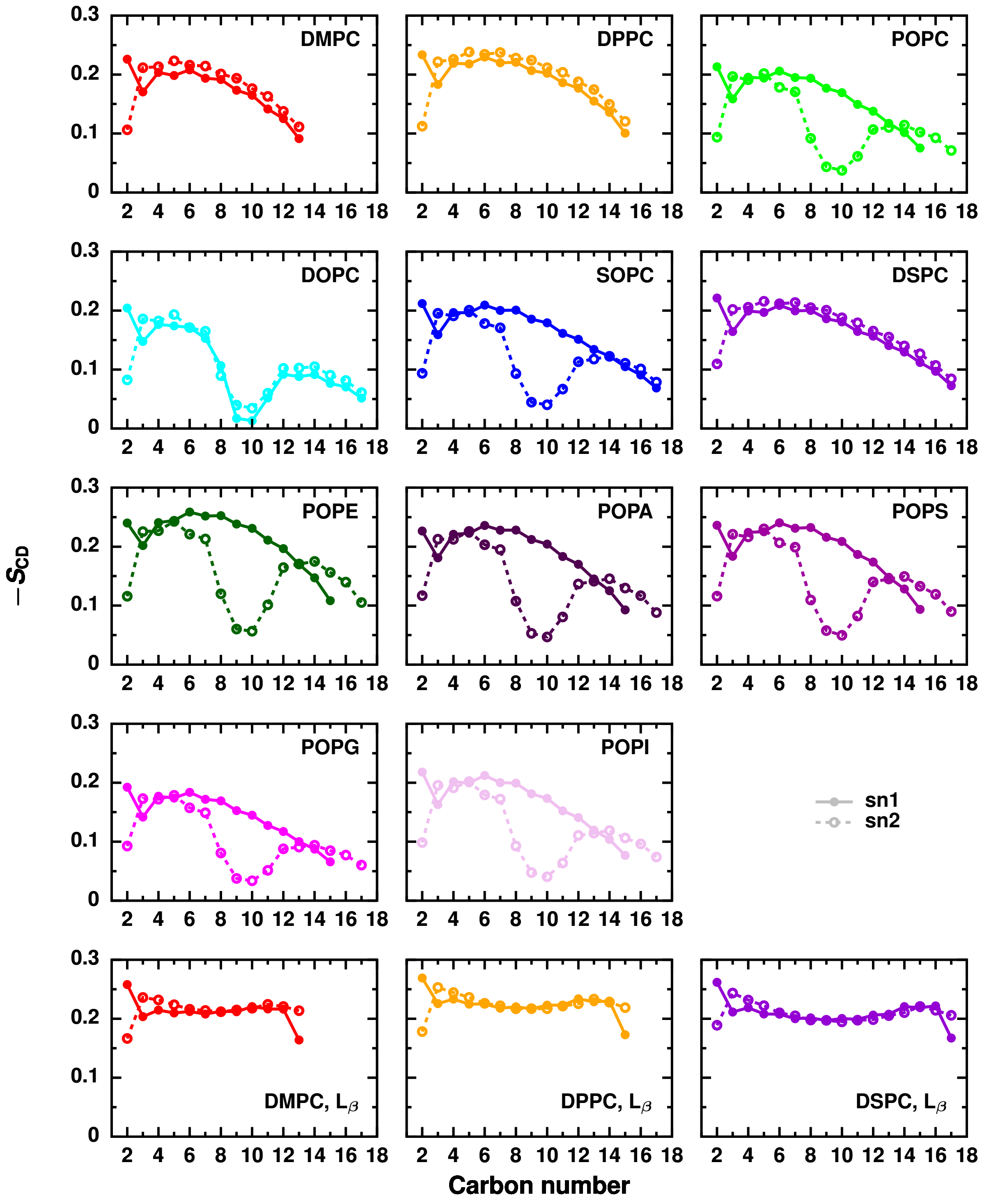


**Figure S2. Lipid chemistry and membrane phase influence the membrane's order.** Deuterium order parameters of the carbons in both acyl chains of the lipids in each membrane.

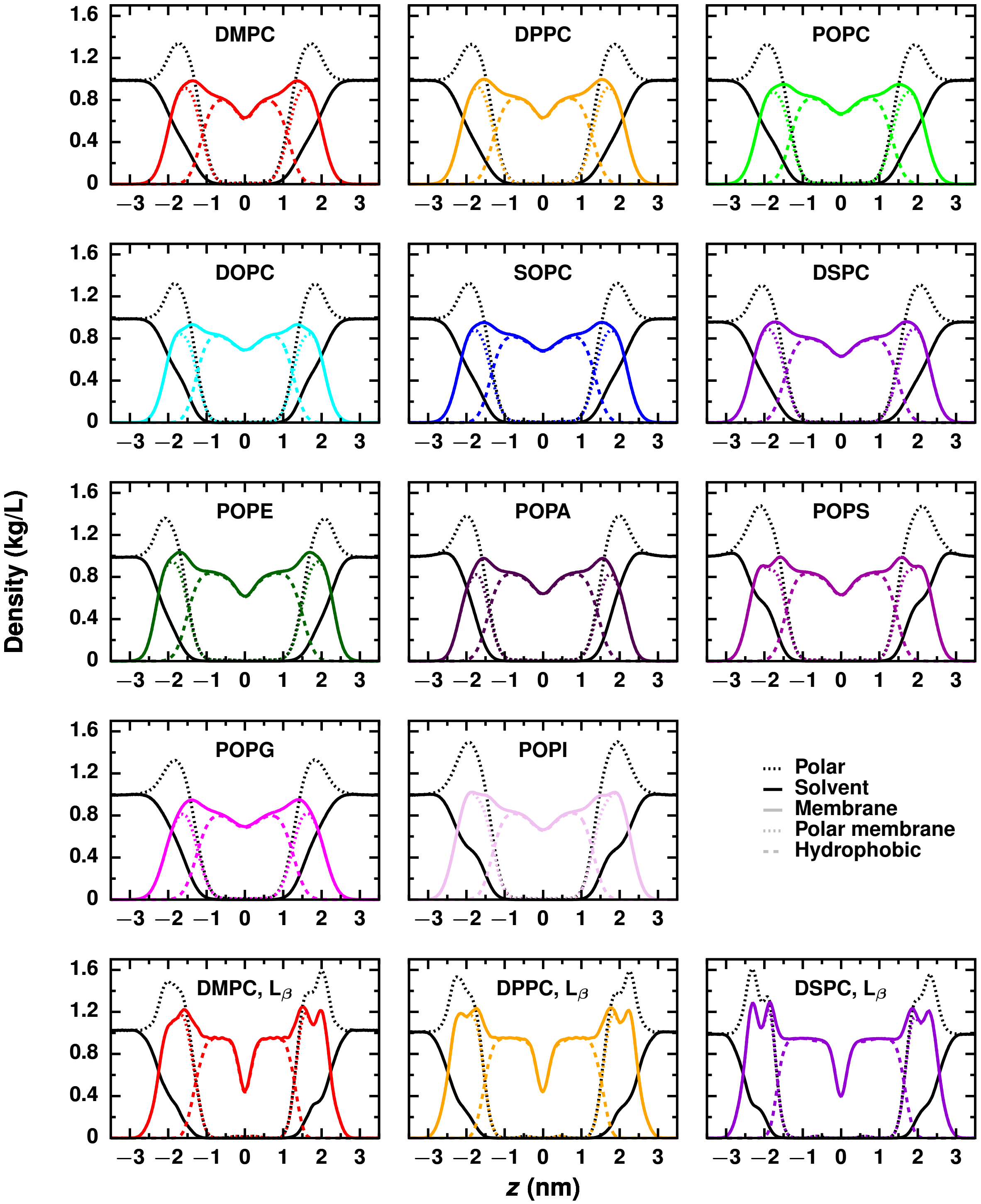


**Figure S3. System composition along the axis normal to the membranes varies with lipid chemistry and membrane phase.** Density profiles along the membrane normal, $z$, for each membrane. The system is grouped into (1) polar, which includes all atoms that are not hydrophobic carbons; (2) solvent; (3) membrane; (4) polar membrane, which includes all lipid atoms that are not hydrophobic carbons; and (5) hydrophobic, which includes all hydrophobic carbons.

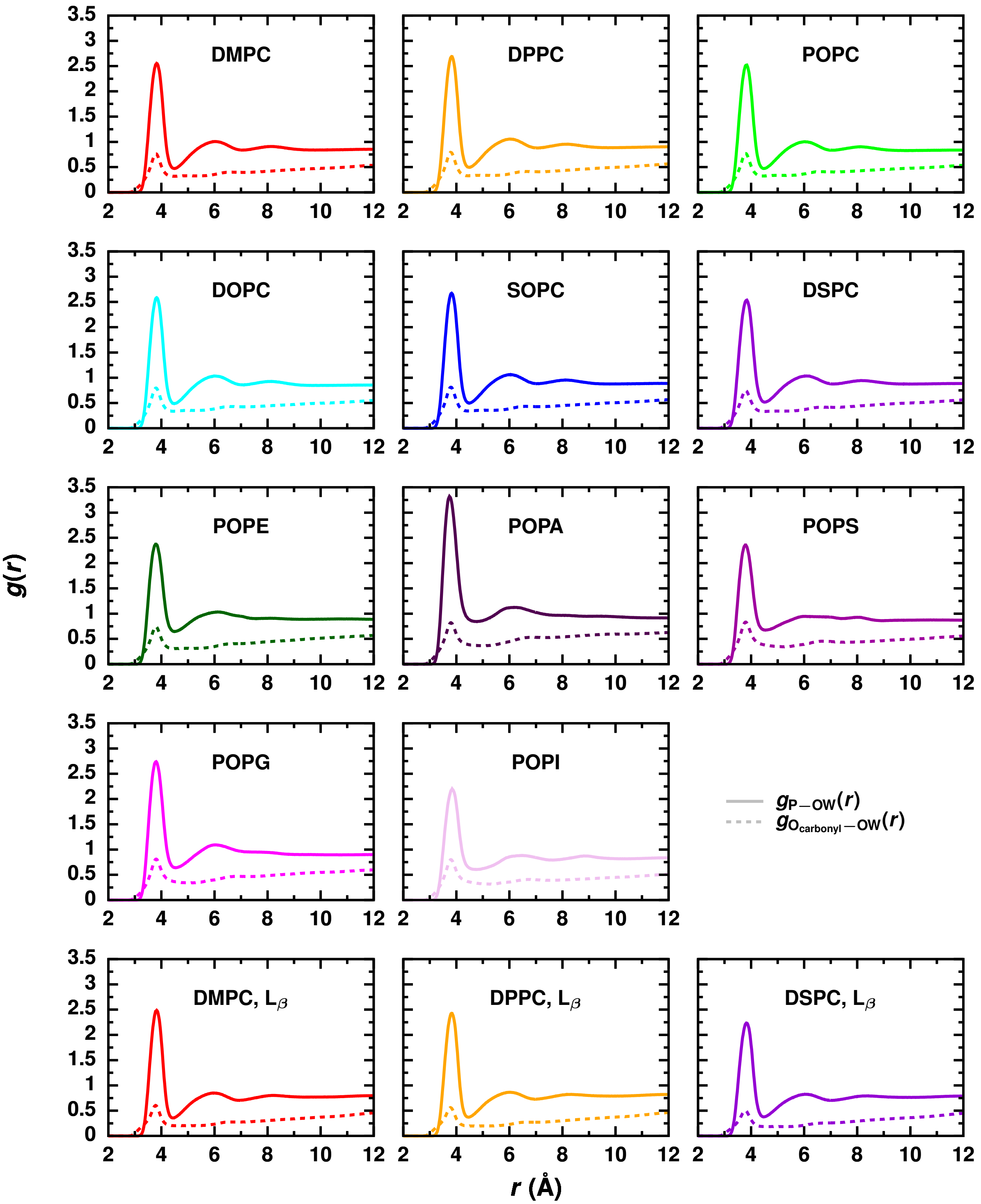


**Figure S4. Water's structure at the membrane surface depends on the lipid headgroup.** Radial distribution functions, $g(r)$, between water oxygen atoms (OW) and either the phosphorous atoms (P) or carbonyl oxygen atoms ($O_{carbonyl}$) of the lipids for each membrane.

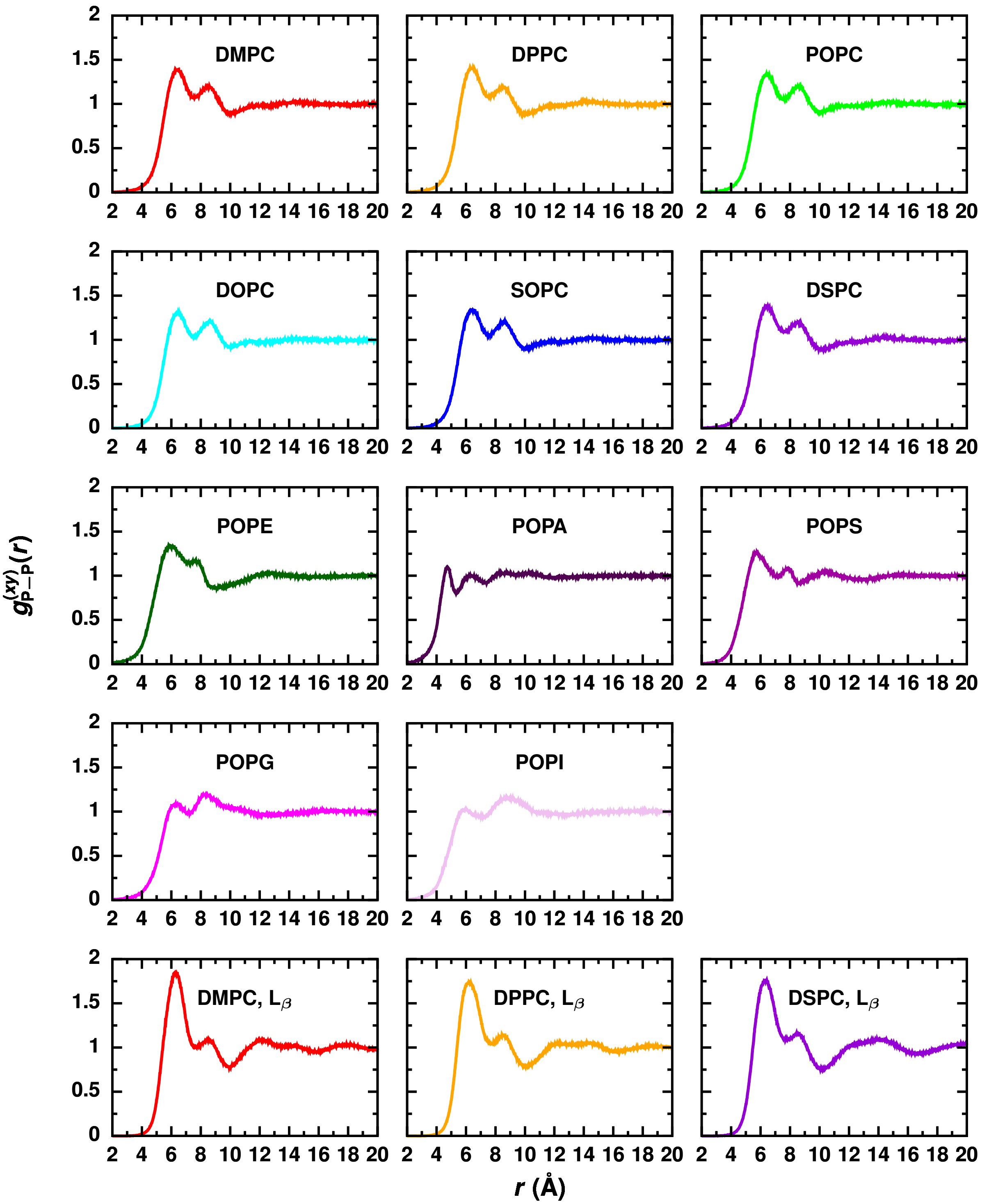


**Figure S5. Membrane surface structure, specifically the arrangement of phosphorous atoms, depends on the lipid headgroup. In $L_\beta$ phase membranes, phosphorous atoms have an increased density of nearest neighbor phosphorous atoms compared to $L_\alpha$ phase membranes.** 2D Radial distribution function, $g^{(xy)}_{P-P}(r)$, between phosphorous atoms of lipids in the same leaflet for each membrane.

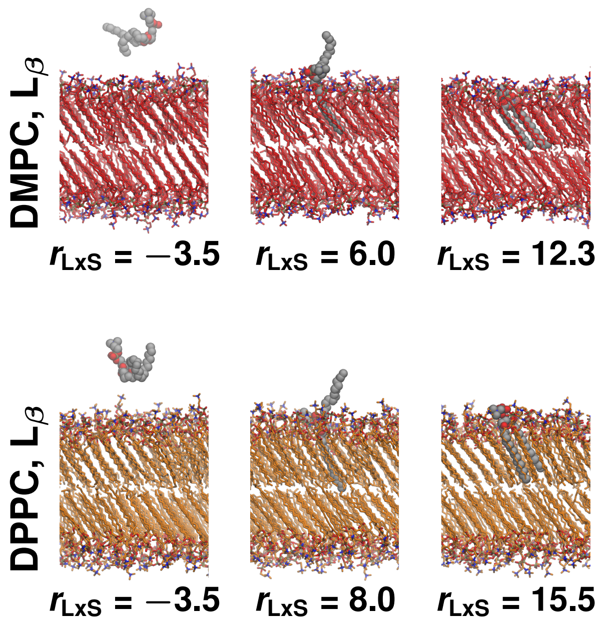


**Figure S6. Splayed lipid configurations are locally thermodynamically stable in $L_\beta$ phase membranes.** Configurations of $L_\beta$ phase (top) DMPC and (bottom) DPPC that are representative of all minimum in $\Delta F(r_{\mathrm{LxS}})$: the lipid in solution, splayed lipid intermdiate, and lipid in the membrane.

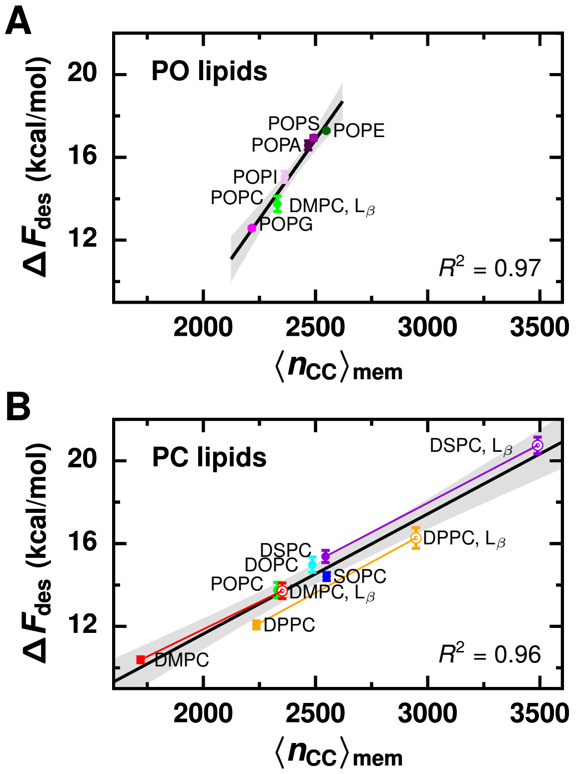


**Figure S7. Desorption barrier is determined by a lipid's local hydrophobic environment, which is measured by the average number of close hydrophobic contacts between membrane lipids, $\langle n_{\rm CC}\rangle_{\rm mem}$.** (A) Plot of $\Delta F_{\rm des}$ versus $\langle n_{\rm CC}\rangle_{\rm mem}$ for lipids with different headgroups. Black line is the best linear fit: $\Delta F_{\rm des} = 0.015\langle n_{\rm CC}\rangle_{\rm mem} - 21.5$. (B) Plot of $\Delta F_{\rm des}$ versus $\langle n_{\rm CC}\rangle_{\rm mem}$ for lipids with different tail chemistry and membrane phase. Colored lines connect points for membranes of the same lipid species but different phase. Black line is the best linear fit: $\Delta F_{\rm des} = 0.0058\langle n_{\rm CC}\rangle_{\rm mem} + 0.05$. Gray region indicates 95% confidence interval.

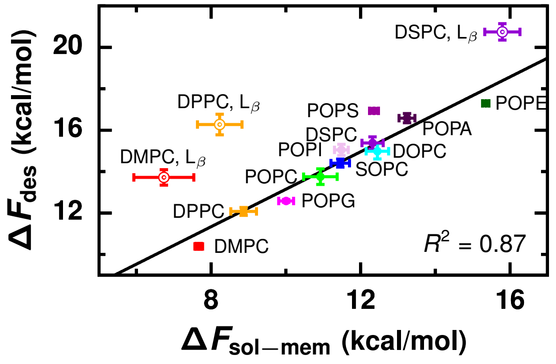


**Figure S8. Desorption barrier is roughly linearly correlated with the transfer free energy, $\Delta F_{\rm sol-mem} = \Delta F(r_{\rm LxS} = -3.45)$** Black line is the best linear fit obtained from weighted least squares, with points weighted by the error in $\Delta F_{\rm sol-mem}$: $\Delta F_{\rm des} = 0.90\Delta F_{\rm sol-mem} + 4.1$.

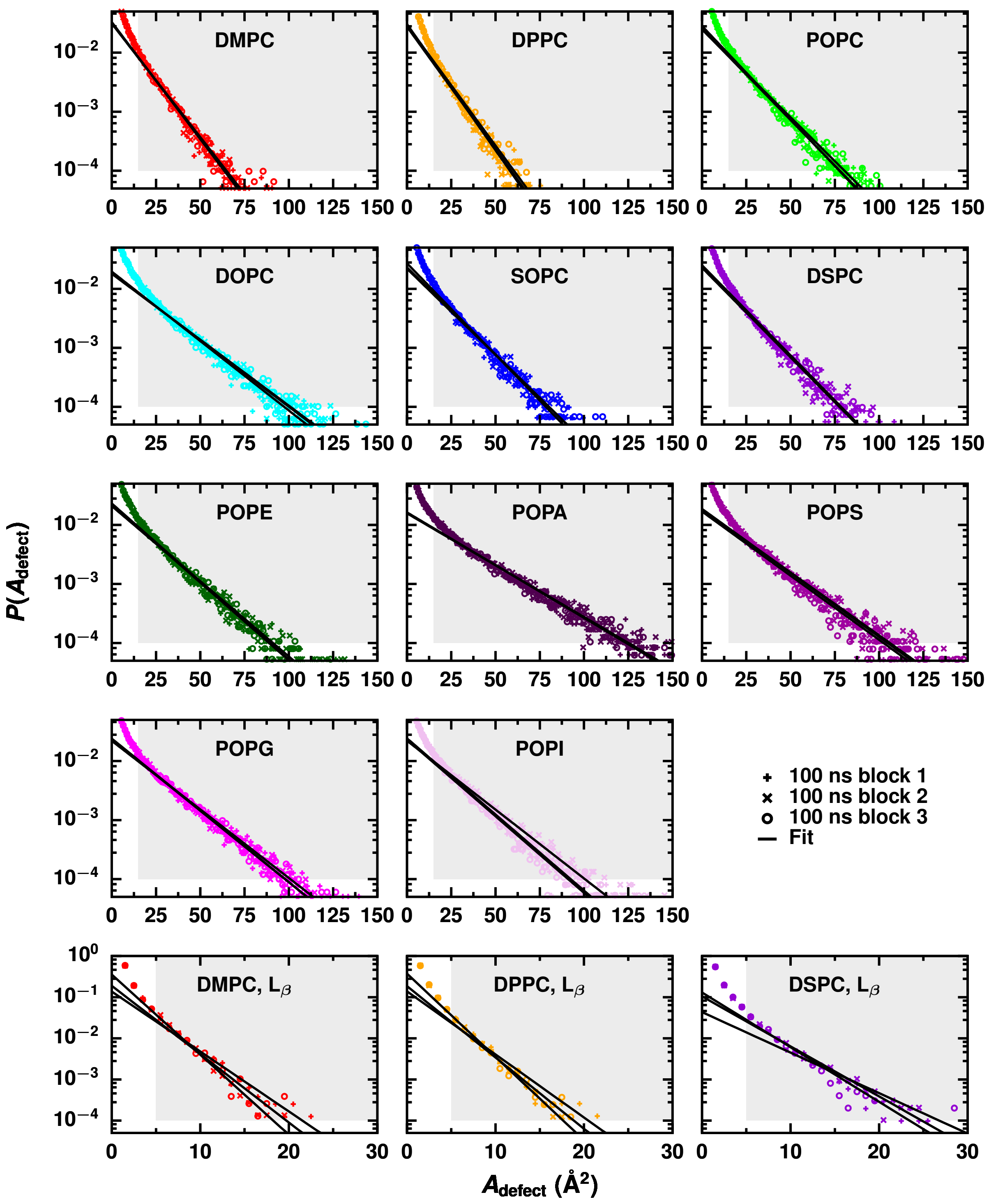


**Figure S9. The size of packing defects that expose the membrane's hydrophobic core to solvent vary with lipid chemistry and membrane phase.** Distributions of the area of packing defects for each membrane. For each membrane, three independent fits were performed to distributions constructed from three 100ns simulation blocks. The region used to fit to an exponential distribution is highlighted in gray.

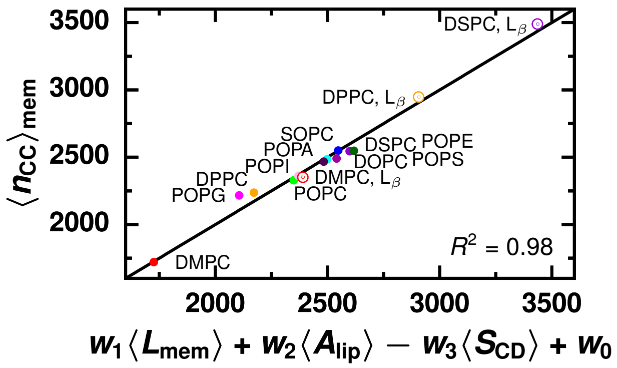


**Figure S10.** $\langle n_{\text{CC}} \rangle_{\text{mem}}$ **varies with membrane thickness, lipid packing, and membrane order.** A linear combination of a membrane's average thickness, $\langle L_{\text{mem}} \rangle$, average area per lipid, $\langle A_{\text{lip}} \rangle$, and average deuterium order parameter of all carbons in the lipid tails, $-\langle S_{\text{CD}} \rangle$, with coefficients $w_1 = 126$ Å$^{-1}$, $w_2 = -40$ Å$^{-2}$, $w_3 = 9964$, and $w_0 = 1584$ accurately models $\langle n_{\text{CC}} \rangle_{\text{mem}}$. Lipids in thicker, more tightly packed (smaller area per lipid), and more ordered (larger $-\langle S_{\text{CD}} \rangle$) membranes form more hydrophobic contacts.

#### SUPPLEMENTAL TABLES

**Table S1. Umbrella sampling parameters.**

| Membrane | Spring constant (kJ/mol) | Location of first center | Location of last center | Number of windows |
|---|---|---|---|---|
| DMPC | 20 | −5.4468 | 13 | 52 |
| DPPC | 40 | −6 | 14 | 54 |
| POPC | 40 | −6 | 14 | 54 |
| DOPC | 40 | −6 | 19.2828 | 68 |
| SOPC | 40 | −6 | 19.2828 | 68 |
| DSPC | 40 | −6 | 19.2828 | 68 |
| POPE | 40 | −6 | 14 | 54 |
| POPA | 40 | −6 | 14 | 54 |
| POPS | 40 | −6 | 14 | 54 |
| POPG | 40 | −6 | 14 | 54 |
| POPI | 40 | −6 | 14 | 54 |
| DMPC, $L_\beta$ | 40 | −6 | 15.1358 | 67* |
| DPPC, $L_\beta$ | 40 | −6 | 17.7735 | 67† |
| DSPC, $L_\beta$ | 40 | −6 | 19.2828 | 68 |

* Count includes 10 windows centered at −0.9048, −0.5273, −0.1499, 1.7372, 2.8695, 7.3985, 7.7760, 8.1534, and 8.9082, which were added to decrease the spacing between some original windows and improve sampling.

† Count includes 3 windows centered at 9.2830, 9.6604, and 9.5660, which were added to improve sampling around a local maximum.

**Table S2. Free energy barriers for lipid desorption ($\Delta F_{des}$), lipid insertion ($\Delta F_{ins}$), and the creation of splayed lipid configurations ($\Delta F_{splay}$) calculated from simulation. Reported error is the standard error of the free energy barrier estimated from four 8 ns simulation blocks. Experimental estimates of the activation free energy for lipid transport ($\Delta F_{expt}$) are provided when available.**

| Membrane | $\Delta F_{des}$ (kcal/mol) | $\Delta F_{ins}$ (kcal/mol) | $\Delta F_{splay}$ (kcal/mol) | $\Delta F_{expt}$ (kcal/mol) |
|---|---|---|---|---|
| DMPC | $10.4 \pm 0.1$ | $2.7 \pm 0.2$ | — | 20.8 – 24.1 (2–4) |
| DPPC | $12.1 \pm 0.2$ | $3.2 \pm 0.4$ | — | 22.9 (2) |
| POPC | $13.8 \pm 0.4$ | $2.8 \pm 0.6$ | — | 22.9 – 23.7 (2,5,6) |
| DOPC | $15.0 \pm 0.4$ | $2.5 \pm 0.5$ | — | 24.2 (6) |
| SOPC | $14.4 \pm 0.2$ | $3.0 \pm 0.3$ | — | |
| DSPC | $15.4 \pm 0.3$ | $3.1 \pm 0.4$ | — | |
| POPE | $17.3 \pm 0.1$ | $1.9 \pm 0.1$ | — | |
| POPA | $16.6 \pm 0.2$ | $3.3 \pm 0.3$ | — | |
| POPS | $16.9 \pm 0.1$ | $4.6 \pm 0.2$ | — | |
| POPG | $12.6 \pm 0.1$ | $2.6 \pm 0.2$ | — | |
| POPI | $15.1 \pm 0.3$ | $3.6 \pm 0.3$ | — | |
| DMPC, $L_\beta$ | $13.7 \pm 0.4$ | $7.0 \pm 0.9$ | $9.4 \pm 0.4$ | 22.2 – 24.8 (2,3,7) |
| DPPC, $L_\beta$ | $16.3 \pm 0.5$ | $8.0 \pm 0.8$ | $11.4 \pm 0.5$ | 25.6 (7) |
| DSPC, $L_\beta$ | $20.7 \pm 0.4$ | $5.0 \pm 0.6$ | $14.1 \pm 0.4$ | |

**Table S3. Average membrane physical properties. Reported error is the standard error calculated from three 100 ns simulation intervals.**

| Membrane | $\langle A_{lip} \rangle$ (Å$^2$) | Av. Thickness (nm) | $\langle n_{CC} \rangle_{mem}$ | $\pi_{defect}$ (Å$^2$) | $\langle E_{mem-solv} \rangle$ (kcal/mol) |
|---|---|---|---|---|---|
| DMPC | $62.88 \pm 0.06$ | $3.523 \pm 0.003$ | $1720 \pm 2$ | $11.10 \pm 0.08$ | $-11361 \pm 9$ |
| DPPC | $60.9 \pm 0.2$ | $3.95 \pm 0.01$ | $2237 \pm 6$ | $10.4 \pm 0.1$ | $-9414 \pm 16$ |
| POPC | $66.77 \pm 0.01$ | $3.834 \pm 0.001$ | $2329 \pm 1$ | $14.1 \pm 0.2$ | $-11717 \pm 1$ |
| DOPC | $70.14 \pm 0.07$ | $3.815 \pm 0.003$ | $2485 \pm 2$ | $19.0 \pm 0.2$ | $-11993 \pm 12$ |
| SOPC | $66.60 \pm 0.05$ | $4.002 \pm 0.003$ | $2550 \pm 2$ | $14.5 \pm 0.2$ | $-11688 \pm 7$ |
| DSPC | $65.04 \pm 0.02$ | $4.401 \pm 0.001$ | $2544 \pm 1$ | $14.25 \pm 0.02$ | $-10934 \pm 2$ |
| POPE | $58.98 \pm 0.06$ | $4.148 \pm 0.004$ | $2548 \pm 2$ | $16.6 \pm 0.2$ | $-11168 \pm 18$ |
| POPA | $62.0 \pm 0.1$ | $3.982 \pm 0.005$ | $2467 \pm 3$ | $24.5 \pm 0.1$ | $-14616 \pm 5$ |
| POPS | $61.62 \pm 0.06$ | $4.042 \pm 0.004$ | $2491 \pm 3$ | $20.1 \pm 0.1$ | $-19022 \pm 53$ |
| POPG | $70.70 \pm 0.07$ | $3.630 \pm 0.003$ | $2216 \pm 2$ | $18.2 \pm 0.2$ | $-13889 \pm 3$ |
| POPI | $65.9 \pm 0.2$ | $3.848 \pm 0.009$ | $2363 \pm 5$ | $17.4 \pm 0.5$ | $-15181 \pm 25$ |
| DMPC, $L_\beta$ | $49.3 \pm 0.1$ | $3.92 \pm 0.05$ | $2351 \pm 5$ | $2.6 \pm 0.2$ | $-9986 \pm 52$ |
| DPPC, $L_\beta$ | $48.75 \pm 0.02$ | $4.376 \pm 0.002$ | $2947 \pm 2$ | $2.5 \pm 0.2$ | $-9416 \pm 16$ |
| DSPC, $L_\beta$ | $50.08 \pm 0.07$ | $4.718 \pm 0.003$ | $3489 \pm 8$ | $3.8 \pm 0.3$ | $-9037 \pm 11$ |

#### SUPPLEMENTAL REFERENCES


1. Pluhackova, K., S. A. Kirsch, J. Han, L. P. Sun, Z. Y. Jiang, T. Unruh, and R. A. Bockmann, 2016. A critical comparison of biomembrane force fields: Structure and dynamics of model DMPC, POPC, and POPE bilayers. *J. Phys. Chem. B* 120:3888–3903.

2. McLean, L. R., and M. C. Phillips, 1984. Kinetics of phosphatidylcholine and lysophosphatidylcholine exchange between unilamellar vesicles. *Biochemistry* 23:4624–4630.

3. Wimley, W. C., and T. E. Thompson, 1990. Exchange and flip-flop of dimyristoylphosphatidylcholine in liquid-crystalline, gel, and 2-component, 2-phase large unilamellar vesicles. *Biochemistry* 29:1296–1303.

4. Nakano, M., M. Fukuda, T. Kudo, H. Endo, and T. Handa, 2007. Determination of interbilayer and transbilayer lipid transfers by time-resolved small-angle neutron scattering. *Phys. Rev. Lett.* 98:238101.

5. Jones, J. D., and T. E. Thompson, 1990. Mechanism of spontaneous, concentration-dependent phospholipid transfer between bilayers. *Biochemistry* 29:1593–1600.

6. Pownall, H. J., D. L. M. Bick, and J. B. Massey, 1991. Spontaneous phospholipid transfer: Development of a quantitative model. *Biochemistry* 30:5696–5700.

7. Xia, Y., M. Li, K. Charubin, Y. Liu, F. A. Heberle, J. Katsaras, B. X. Jing, Y. X. Zhu, and M. P. Nieh, 2015. Effects of nanoparticle morphology and acyl chain length on spontaneous lipid transfer rates. *Langmuir* 31:12920–12928.
